# Exploration of phosphoproteomic association during epimorphic regeneration

**DOI:** 10.1101/2024.03.08.584197

**Authors:** Sarena Banu, P V Anusha, Komal Mandal, Mohammed M Idris

**Affiliations:** CSIR-CCMB, Uppal Road, Hyderabad, India

**Keywords:** Regeneration, Proteomics, Phosphoprotein, TiO2, Immunoprecipitation, Zebrafish

## Abstract

Unravelling the intricate patterns of site-specific protein phosphorylation during Epimorphic regeneration holds the key to unlocking the secrets of tissue complexity. Understanding these precise modifications and their impact on protein function could shed light on the remarkable regenerative capacity of tissues, with potential implications for therapeutic interventions. In this study we have systematically mapped the global phosphorylation modifications within regenerating tissue of zebrafish caudal fins, elucidating the intricate landscape of signalling pathway associate with the regeneration process. A total of 74 and 440 proteins were found undergoing differentially phosphorylated during the process of regeneration from 12hpa to 7dpa against control based on TiO2 column enrichment and immuno precipitation using phosphoserine, phosphothreonine and phosphotyrosine antibodies respectively. Interestingly 95% of the proteins identified from TiO2 enrichment method were also found to be identified through the phosphoprotein antibody pull down method impacting the high accuracy and significance of the methods and greater association of the 70 proteins undergoing differential phosphorylation during the process of regeneration. Whole mount immunohistochemistry analysis reveals high association of phosphorylation at 1dpa, 2dpa and 3 dpa regeneration time points. Based on network pathway analysis it was evident that Fc Receptor-mediated Phagocytosis in Macrophages and Monocytes, Actin cytoskeleton signaling, HGF signaling and Insulin receptor signaling are the most highly associated network pathways for regeneration through differential phosphorylation. This research enhances our comprehension on protein post-translational modification in the context of zebrafish caudal fin tissue regeneration, shedding light on its prospective application in the field of regenerative medicine.

## Introduction

Regeneration is the innate ability of living organisms to replace worn-out parts, repair damaged organs, or reconstitute the entire body from a single cell into a multifaceted organism involving growth, morphogenesis, and differentiation^1^. The regeneration capacity varies significantly between taxonomic groups, for example, Urodele amphibians and Teleost fish demonstrate remarkable efficiency in regenerating intricate structures and organs like limbs, hearts, and kidneys, whereas mammals lack the ability to regenerate any of these complex structures^2,3^. The regenerative capacity allows us to explore critical factors that influence the development of tissues and organs, generating mechanistic insights into the developmental basis of regeneration in higher species.

Among the few vertebrate taxa, zebrafish are the most exceptional organisms capable of reinstating a wide range of tissues and organs, including their locomotive appendages^3^. The zebrafish appendage/caudal fin, exhibits lower complexity and rapid regeneration, making it a potent model for epimorphic regeneration^3,4^. Zebrafish caudal fin amputation activates proximal epidermal cell migration from the stump margins to shield the exposed stump by establishing wound epithelium within 12 hours-post-amputation. In the next 1-2-day post amputation (dpa), the wound epithelium stiffens forming an apical epidermal cap (AEC), followed by rehabilitation of extracellular matrix and pre-programmed accumulation of morphologically homogenous clump of blastema at the amputation plane. Subsequently, between 3-7 dpa, skeletal re-patterning, including outgrowth and bifurcation of new bony fin rays is mediated by the onset of progressive dedifferentiation, proliferation and redifferentiation of mature osteoblasts which continues until the original tissue architecture is reconstituted. By the end of 10 dpa, the fin regenerates completely having original structure and function^3,5^.

Significant progress has been made in understanding the molecular mechanisms underlying regeneration. A single-cell transcriptomic study has mapped the cellular diversity of zebrafish caudal fin regeneration, revealed the enrichment of all three epithelial layers, hematopoietic cells, and mesenchymal cells within the amputated fin region, along with their associated mechanisms^6^. Our recent study further elucidated the regeneration mechanism by mapping the differential gene expression of 1408 genes and 661 proteins, along with their associated pathways, including cell cycle activation, oxidative phosphorylation, DNA damage response pathways, and many other pathways in zebrafish caudal fin regeneration tissues^5^. It is well known that codon biases lead to active gene turnover into proteins, which subsequently undergo post-translational modifications to become functional. Our previous global proteomics study mapped numerous proteins and their associated pathways related to cytoskeleton remodelling and the immune response^7^. The 2D-based proteomic analysis revealed significant expression levels of annexin, a calcium-binding protein which undergoes continuous phosphorylation modifications^7,8^. A subsequent study demonstrated that the knockout of Annexin family genes (Annexin a2a and Annexin a2b) significantly represses other annexin family proteins and effectively halts the regeneration process of the caudal fin^9^. Phosphorylation represents the most prevalent and extensively studied regulatory post-translational modification within biological systems^10,11^. In zebrafish, protein tyrosine phosphatase have been identified during caudal fin regeneration^12^. Remarkably, despite its significance, no study has been conducted to investigate protein phosphorylation modifications in zebrafish caudal fin regeneration.

Phosphoproteomics has emerged as a powerful tool for identifying protein phosphorylation sites both qualitatively and quantitatively^11,13^. Understanding site-specific phosphorylation is crucial for unraveling fundamental biological processes. In vertebrates, the primary focus of research has been on the phosphorylation of hydroxyl-containing amino acid residues, namely serine, threonine, and tyrosine due to their abundance. Protein phosphorylation, orchestrated by protein kinases and counterbalanced by protein phosphatases, is linked with additional regulatory mechanisms such as second messengers and protein-protein interactions (PPIs). This collective orchestration extends to enzymes like GTPases, phosphatases, and lipid kinases, collectively governing the intracellular signaling dynamics of cellular functions. Furthermore, protein phosphorylation is a critical post-translational modification (PTM) present across various life forms, and its dysregulation in human cells is closely linked to diseases such as cancer and diabetes^14^. Thus, global phosphoproteomics aids in understanding the critical phosphorylation events and signaling pathways that govern the cellular process. Our study has mapped the global phosphorylation modifications of zebrafish caudal fin regenerating tissues and the associated signaling pathways. This research contributes to our understanding of protein post-translational modification in zebrafish caudal fin tissue regeneration and its potential applications in regenerative medicine.

## Materials and Methods

### Animal housing and Regeneration experiment

Adult wild type zebrafish (6-12 months old) were collected from the CCMB zebrafish laboratory and maintained on a 14-hour light/10-hour dark cycle. Water temperature and pH are maintained under standard laboratory conditions. The zebrafish were anaesthetized with 0.1% tricaine and amputated on the caudal fin distal region for regeneration experiments, as previously described^8^. The regenerated caudal fins were collected at 0hpa (hours post amputation), 12hpa, 1dpa (day post amputation), 2dpa, 3dpa, and 7dpa. The experimental protocol was approved by Centre for Cellular and Molecular Biology institutional animal ethics committee (IAEC/CCMB/Protocol #50/2013).

### Phosphopeptide Enrichment using TiO2 column

Total protein was extracted from zebrafish fin tissue (5 animals per batch) at each regenerating time point (0hpa,12hpa,1dpa,2dpa,3dpa,7dpa) and quantified using the amido black assay, as previously described^5,8^. 50μg protein was loaded on 10% SDS PAGE, sliced, and digested with trypsin^5,8,15^. The digested peptides were desalted before TiO2 enrichment. Vacuum dried desalted peptides were dissolved in binding buffer, loaded on an equilibrated TiO2 column, washed, and eluted according to the manufacturer’s protocol (High-SelectTM TiO2 Phosphopeptide Enrichment Kit, Thermo Fisher). The enriched phosphopeptides were dissolved in 0.1% formic acid for Liquid Chromatography Mass Spectrometry (LCMS/MSMS) analysis^15^. The obtained raw data were analysed using the PTM Analysis Workflow with the ptmRS Node in proteome discoverer 2.2, and peptides were quantified using label free quantification.

### Immunoprecipitation (IP)

The total protein was extracted from 15 animals fin tissue for each time points (0hpa,12hpa,1dpa,2dpa,3dpa,7dpa) using IP lysis buffer (20 mM Tris; adjust to pH 7.4 5 mM EDTA,1mM NaF,150mM NaCl, 1mM sodium orthovanadate, 1% Triton X-100, Protease Inhibitor tablet, Aprotinin, leupeptin and pepstatin). The samples were homogenized, continuously agitated on an orbital shaker at 4 °C for 2 hours, and then centrifuged for 30 minutes at 14000 rpm to remove tissue debris. The extracted total protein was quantified using the amido black assay, as previously described^8^. For all regenerating time points, immunoprecipitation was performed with 200μg of protein, 5ul of phosphoserine, phosphothreonine and phosphotyrosine antibodies (Merck Millipore, USA). The protein and antibody mixture were incubated at 4°C overnight. The next day, prewashed Protein G agarose beads (Millipore) were added to the protein antibody mix and incubated at 4°C for 4 hours on an orbital shaker. This lysate-beads mixture was centrifuged at 2000 RCF and washed three times in IP wash buffer (20 mM Tris; adjust to pH 7.4, 5 mM EDTA,1mM NaF,150mM NaCl, 1mM sodium orthovanadate, Protease Inhibitor tablet, Aprotinin, leupeptin and pepstatin). Immunoprecipitated proteins were eluted in 2x loading buffer by boiling agarose beads at 100°C for 5 minutes. The proteins were immediately loaded on 10% SDS PAGE, fractionated, trypsin digested, and purified using spin C18 columns. Eluted peptides were loaded into Liquid Chromatography Mass Spectrometry (LCMS/MSMS) with a 90-minute gradient^15^. The obtained raw data were analysed against the *Danio rerio* database in proteome discoverer 2.2 using the PTM Analysis Workflow with the ptmRS Node and filtered with respective phosphorylated amino acids (Serine, Threonine and Tyrosine).

### Whole Mount Immunohistochemistry

Whole mount immunohistochemistry was performed in biological duplicates for all the regenerating time points (0hpa,12hpa,1dpa,2dpa,3dpa, and 7dpa). The amputated caudal fin tissues were fixed in 4% paraformaldehyde, dehydrated and rehydrated with methanol gradients, and permeabilized in acetone as previously described^8^. Fin tissues were incubated in 1:100 dilutions of phosphoserine, phosphothreonine and phosphotyrosine primary antibodies (Merk Millipore, USA) for 48 hours, washed four times with PBS and incubated in respective FITC conjugated anti mouse and anti-rabbit secondary antibodies (1:200) for 2 hours. The fluorescence signal was imaged with a Zess Axio imager Z2 apotome microscope using ZEN 2.6 pro software.

### Western Blot

Total protein was extracted from 8 animal’s fin tissue (over all 48 animals) at each regenerating time points (0hpa,12hpa, 1dpa, 2dpa,3dpa, 7dpa). Total proteins were extracted and quantified as previously described^8^. A Western blot was performed with 50μg of protein, electrophoresed and wet transferred into a PVDF membrane^8^. Phosphorylated serine, threonine, and tyrosine protein signals were detected using 1μg of phosphoserine, phosphothreonine and phosphotyrosine primary antibodies (Merk Millipore, USA) and 1:3000 dilution of respective anti-mouse and anti-rabbit secondary antibodies. ODC antibody (1:100 dilution) was used as a control.

### Network Pathway Analysis

Network pathway analysis was performed on the list of proteins identified from both TiO2 enrichment and antibody pull down assay. The differentially expressed phosphoproteins were analysed for the association in canonical pathways, disease & disorders, molecular & cellular functions and physiological system development & functions using Ingenuity Pathway analysis (IPA) software.

## Results

### Phosphoproteins associated with regeneration

Based on Phosphopeptide enrichment using TiO2 column it was found that a total 74 proteins were found differentially phosphorylated and associated with zebrafish caudal fin regeneration with at least one log fold change in one of the regenerating time points (Supplementary Table 1). Proteins such as uridine-cytidine kinase –like 1 isoform X1 (UCKL1), DDRGK domain containing protein 1 precursor (DDRGK1) and vitellogenin (VTG1) were few of the protein found to be phosphorylated majorly during the process of regeneration. Similarly, proteins such as calcium/calmodulin-dependent protein kinase type II delta 1 chain isoform X6 (CAMK2d1), col6a3 isoform X1 (COL6A3) and disabled homolog 2-interacting protein (DAB2) were found dephosphorylated during the process of regeneration (Figure 1).

**Figure 1:**
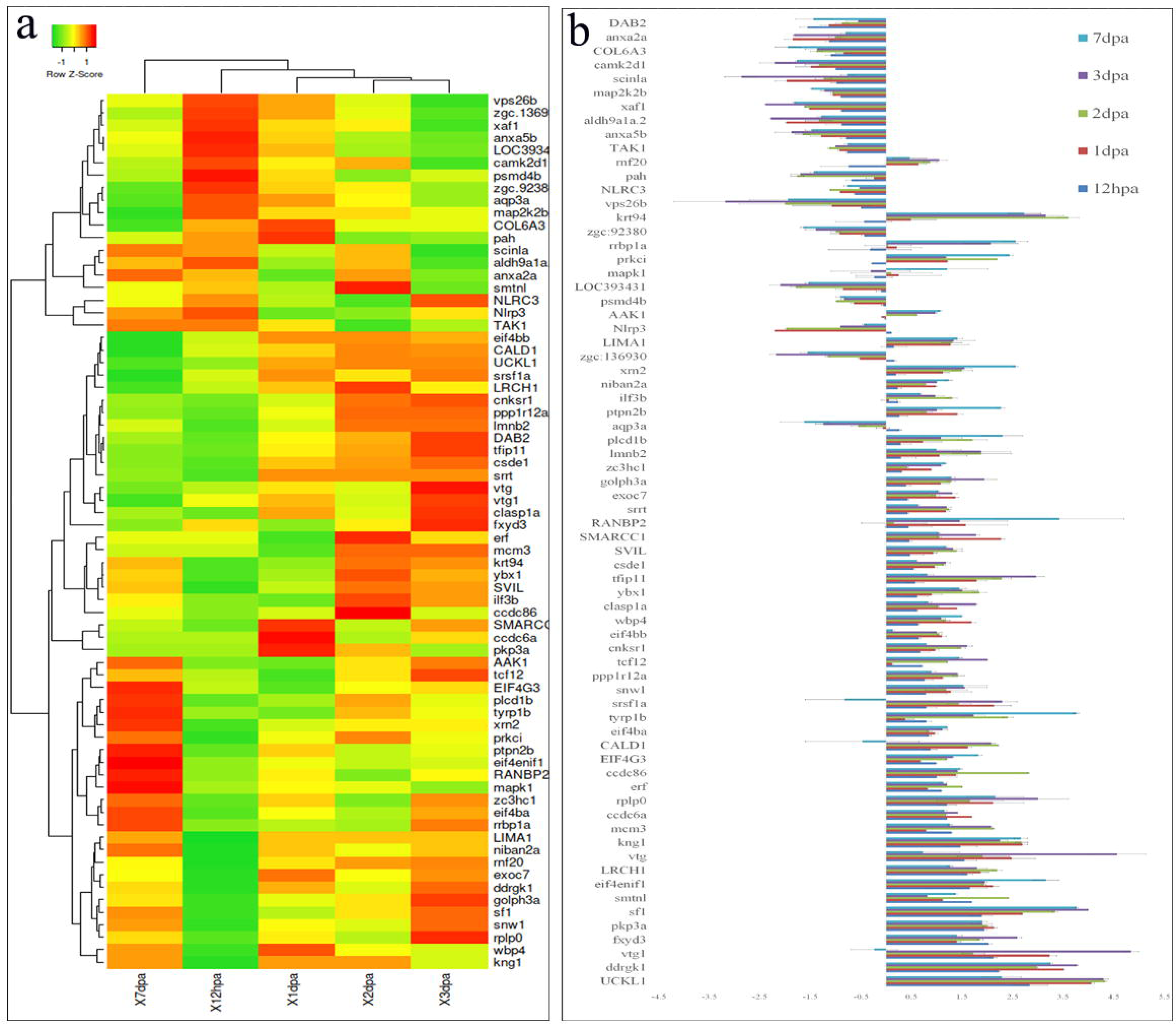
Heat map analysis and Graphical Representation of fold changes of differentially phosphorylated proteins in regenerating zebrafish caudal fin tissue relative to non-regenerating tissue, a. Phosphoproteome heat map analysis and b. Expression pattern analysis of phosphoproteins relative to control represented as the meanL±LSEM.

Based on immunoprecipitation pulldown assay involving three different phosphoproteins antibodies it was found that a total of 440 proteins were found differentially phosphorylated during the process of regeneration (Supplementary Table 2). The list includes 373, 233 and 98 proteins as identified based on phosphoserine, phosphothreonine and phosphotyrosine antibodies respectively. Notably 57 proteins were commonly identified across all three phosphorylated antibody pull downs. Serine/arginine repetitive matrix protein 2 (Srrm2) and non-muscle caldesmon-like (CALD1) were found most phosphorylated and PR domain zinc finger protein 2 (PRDM2) and collagen alpha-2(I) chain precursor (Col1a2) were found dephosphorylated based on immunoprecipitation study.

Interestingly 95% of the proteins (70 proteins) from TiO2 column method were also identified based on antibody pull down method, validating the process of identification. Similarly, the expression pattern of differential phosphorylation was found to be very significant for almost all the 70 proteins and for all the time points (Table 1 and Figure 1). Based on heatmap analysis it was found that phosphoproteomic changes were closely knitted at the 1, 2 and 3 dpa regenerating time points which are out rooted with the wound healing and structural regeneration time points (Figure 1a).

**Table 1:**
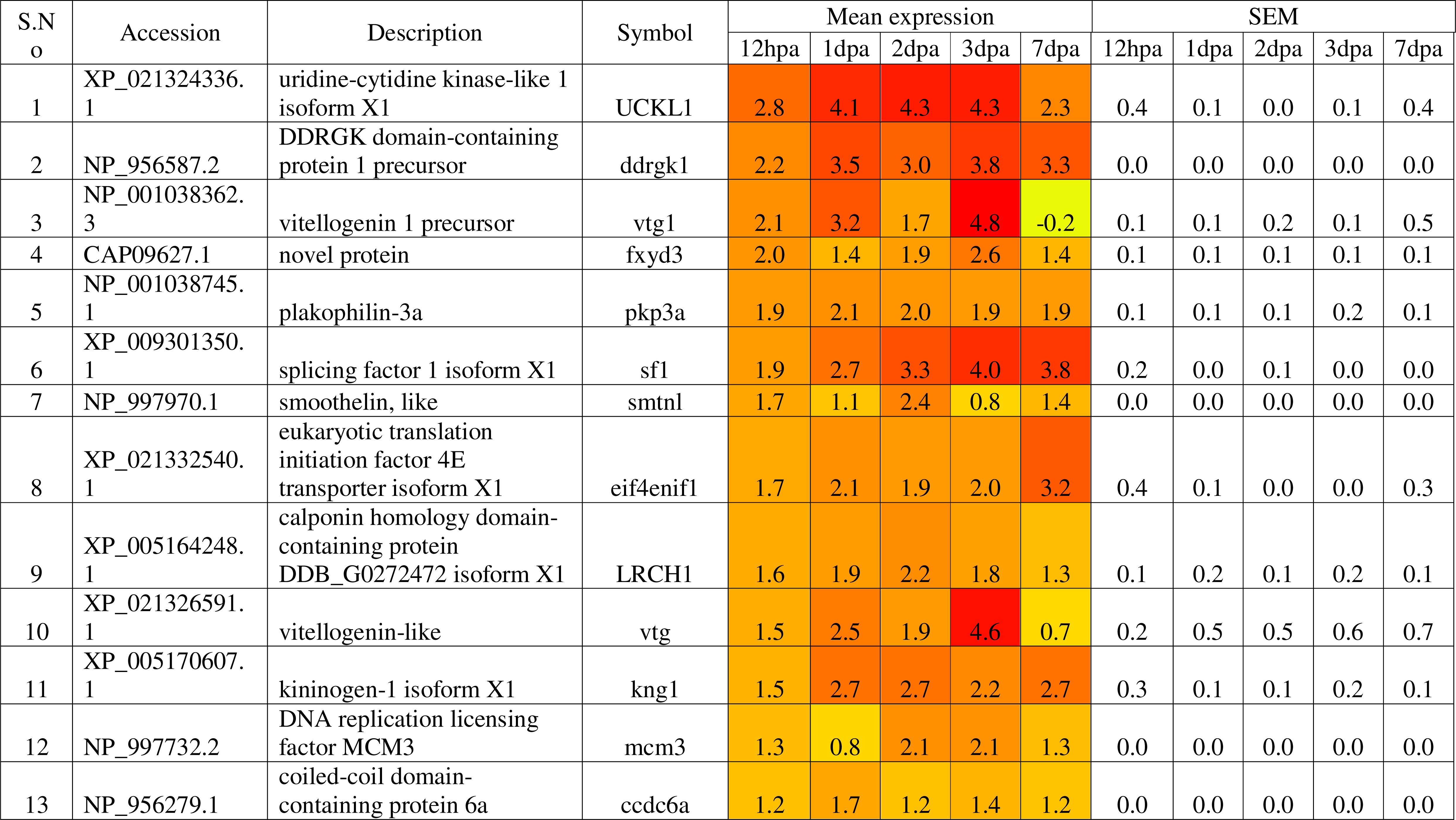
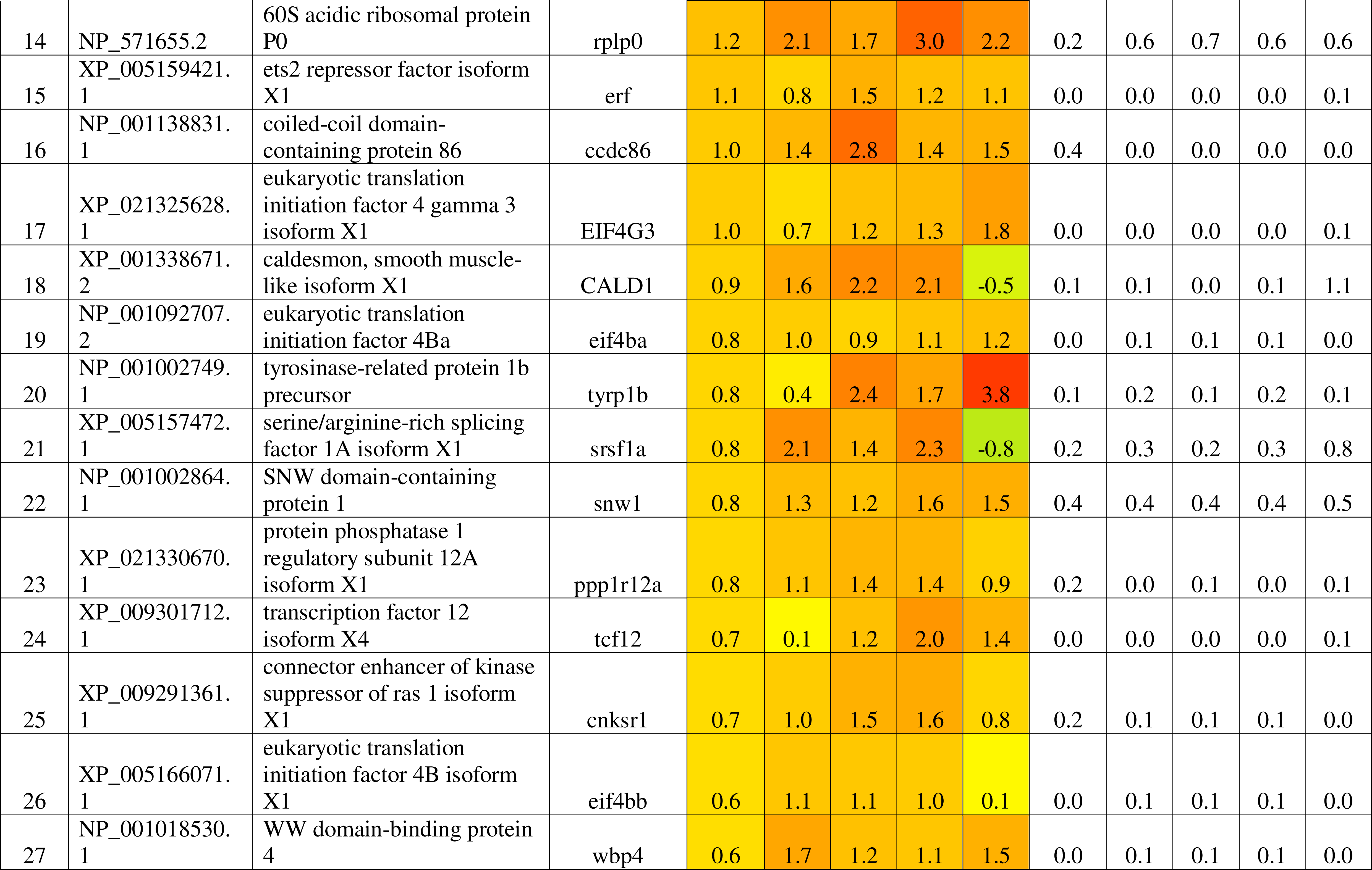
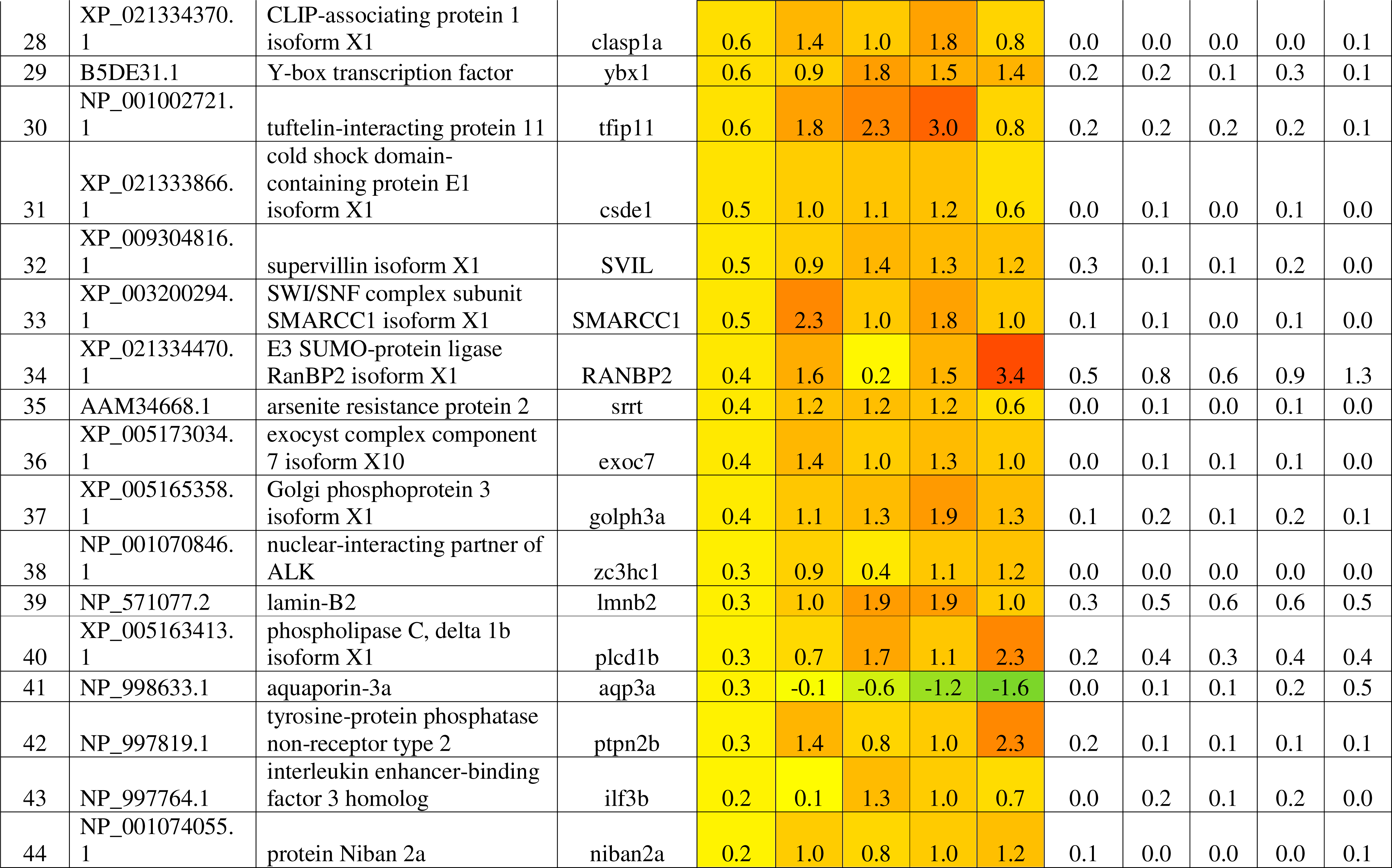
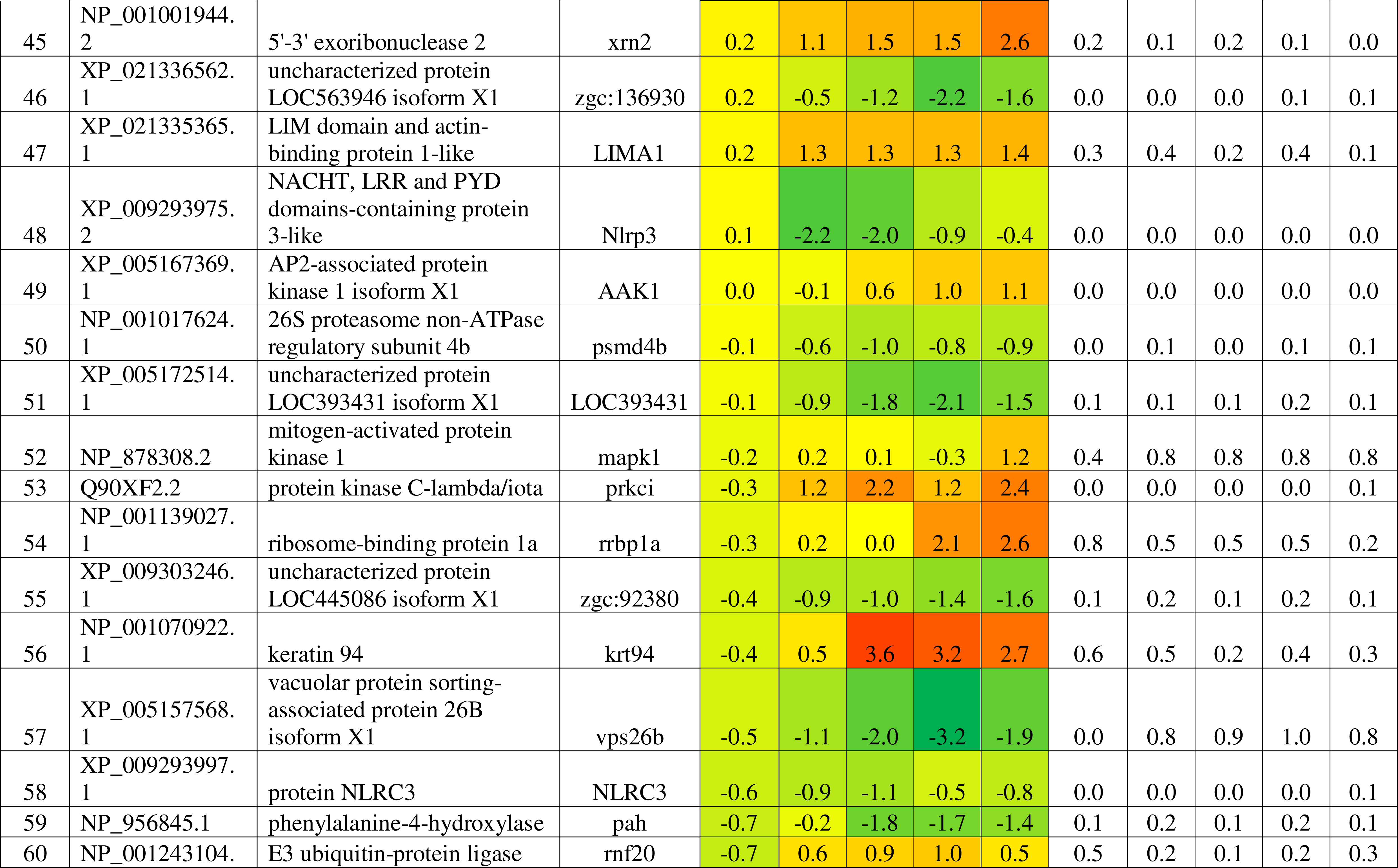
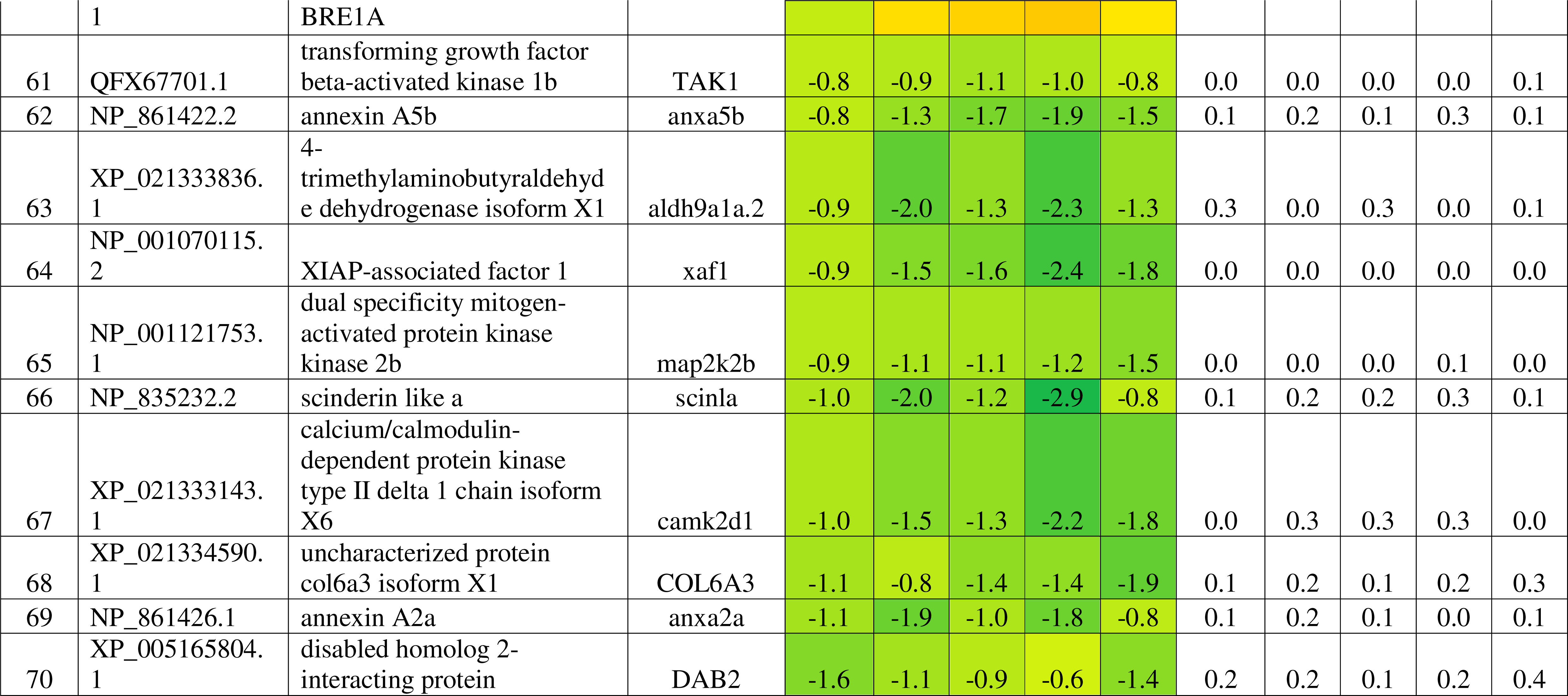
List of phosphoprotein differential expressed during zebrafish caudal fin regeneration based on both TiO2 column enrichment and immunoprecipitation methods.

Based on whole mount immunochemistry analysis it was found that all the three-antibody showed association with phosphorylation in the regenerating caudal fin tissue (Figure 2). Serine and threonine-based phosphorylation was found throughout the regeneration time points with the maximum at 2dpa and 3dpa respectively, whereas tyrosine showed less phosphorylation in comparison to the other antibodies. Western blot analysis showed multiple proteins undergoing phosphorylation for all the three-antibody analysis (Figure 3). As like whole mount immunohistochemistry maximum protein phosphorylation was observed with serine and threonine antibodies.

**Figure 2:**
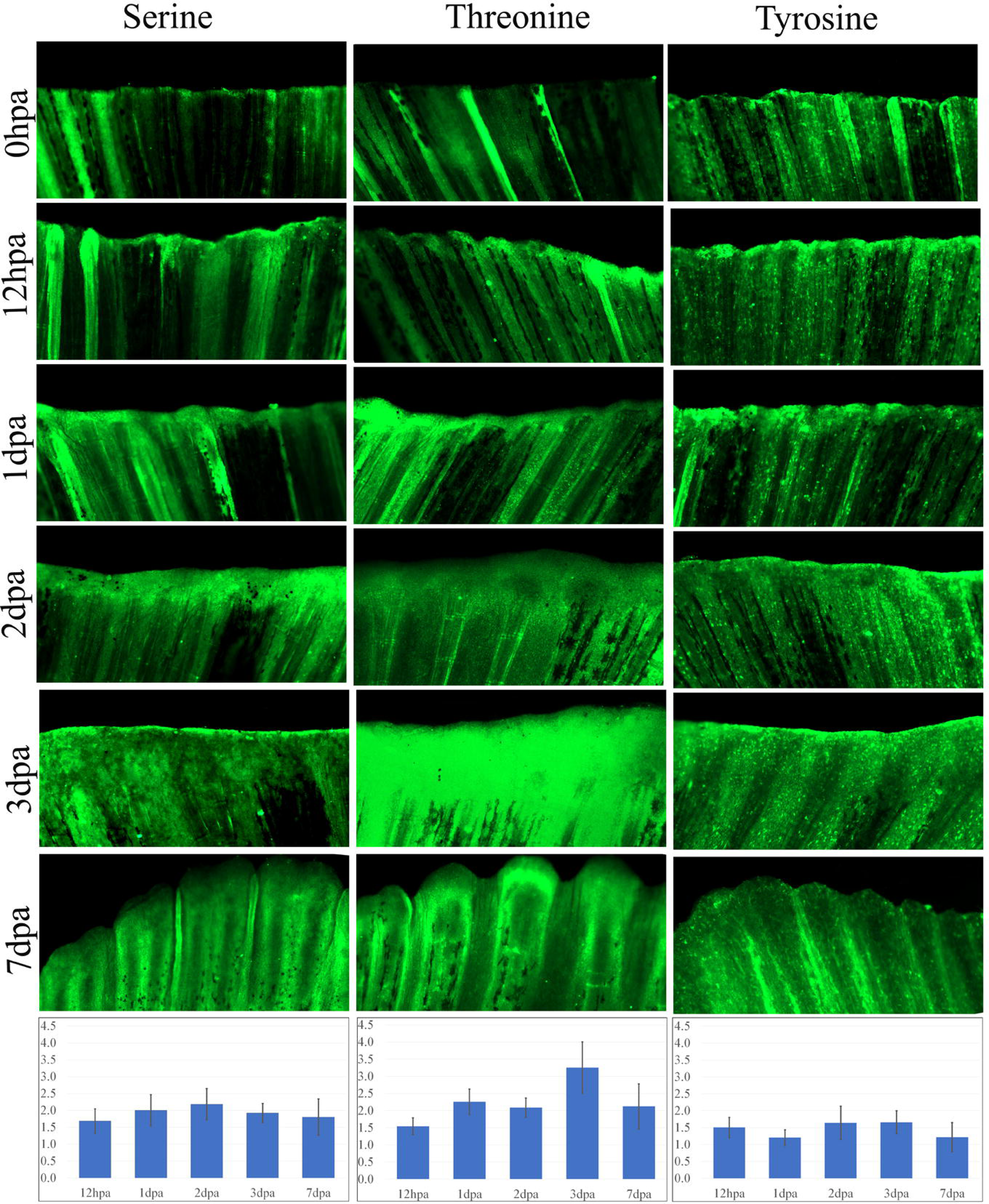
Wholemount immunohistochemistry analysis of differential phosphorylation associated with zebrafish caudal fin regeneration and expression plot at each timepoints with respect to phosphoserine, phosphothreonine and phosphotyrosine antibodies respectively.

**Figure 3:**
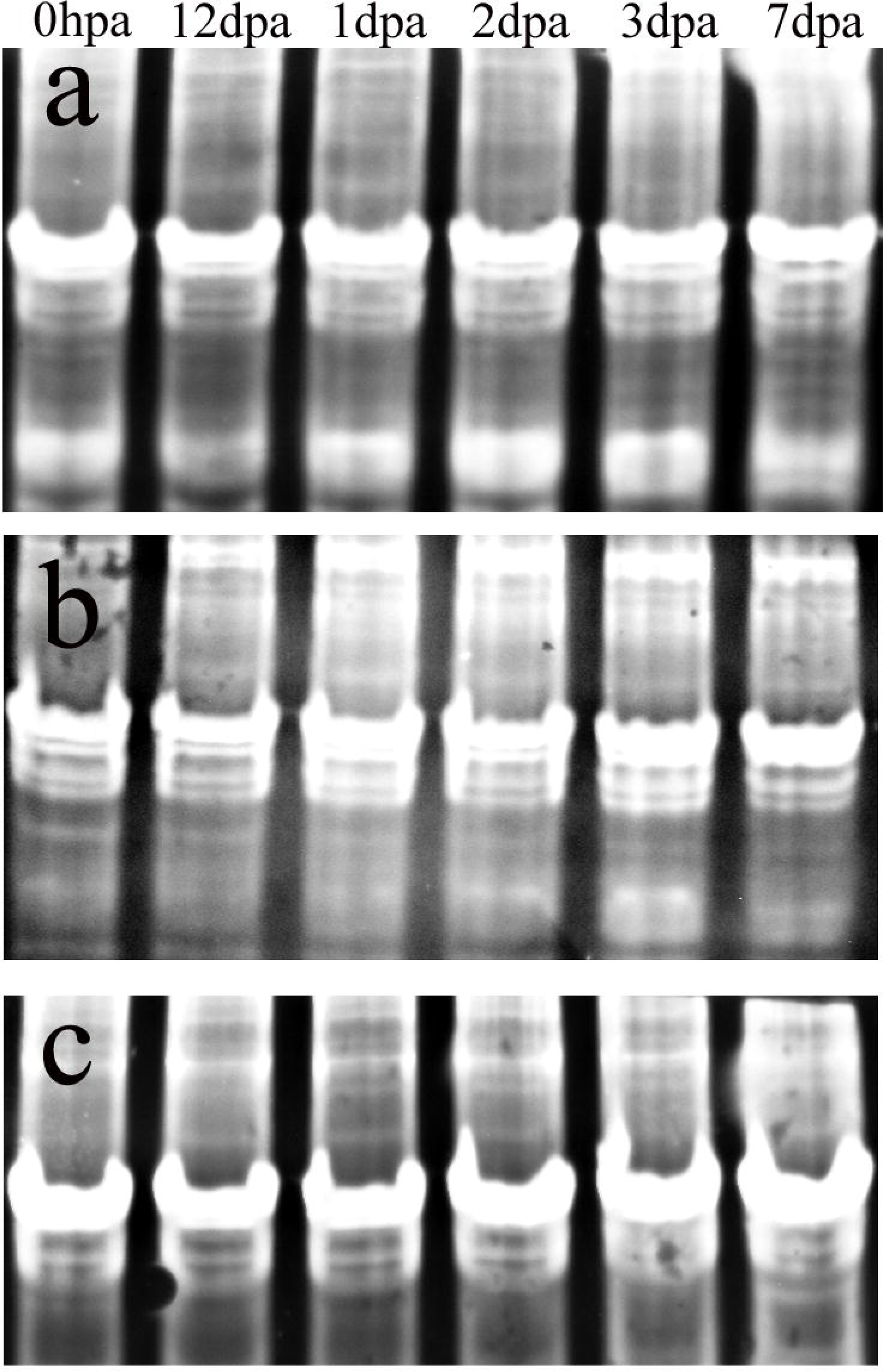
Western blot analysis depicting phosphoproteins for the antibodies at different timepoints 0hpa, 12hpa, 1dpa, 2dpa, 3dpa and 7dpa. a. Blot with phosphoserine antibody, b. Blot with phosphothreonine antibody, and c. Blot with phosphotyrosine antibody.

### Network Pathway analysis

From on the list of proteins identified as undergoing differential phosphorylation for regeneration it was found that Fc Receptor-mediated Phagocytosis in Macrophages and Monocytes, Actin Cytoskeleton Signaling, HGF signaling, Insulin Receptor signaling are the most highly associated canonical pathways (Table 2a). Cancer & organismal injury and abnormalities are the most associated diseases and disorders (Table 2b). The major molecular and cellular functions associated with the regeneration process are cellular assembly and organisation, cellular function and maintenance, cellular movement, cellular development and cellular growth and proliferation (Table 2c). The physiological system development and function associated with regeneration based on phosphorylation are nervous system development and function, tissue development, organismal survival, organismal development, and embryonic development (Table 2d).

**Table 2:**
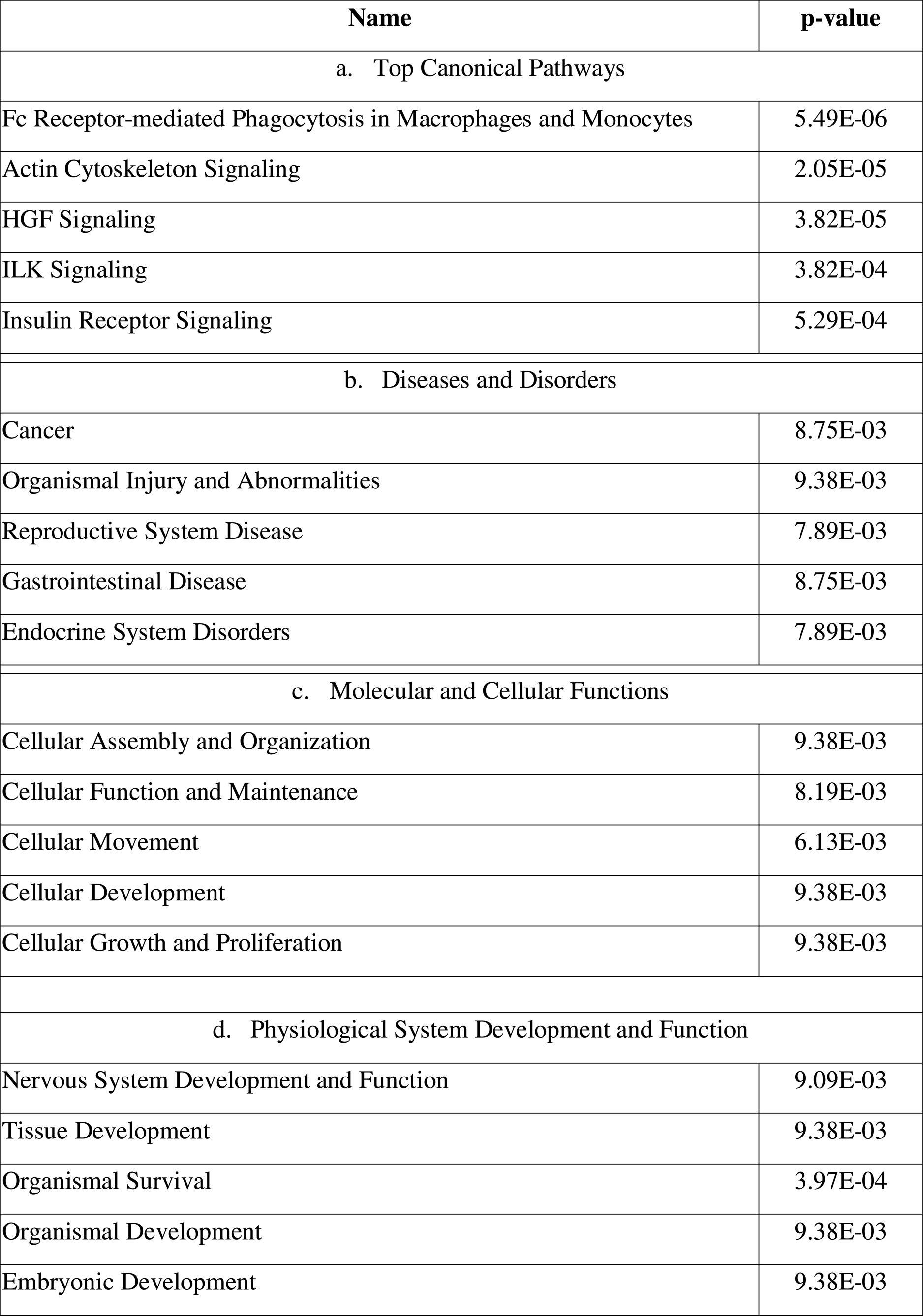
List of top five canonical pathways, Disease and Disorders, Molecular and Cellular functions and d. Physiological System Development and Function.

| Name | p-value |
| --- | --- |
| a. Top Canonical Pathways |  |
| Fc Receptor-mediated Phagocytosis in Macrophages and Monocytes | 5.49E-06 |
| Actin Cytoskeleton Signaling | 2.05E-05 |
| HGF Signaling | 3.82E-05 |
| ILK Signaling | 3.82E-04 |
| Insulin Receptor Signaling | 5.29E-04 |
| b. Diseases and Disorders |  |
| Cancer | 8.75E-03 |
| Organismal Injury and Abnormalities | 9.38E-03 |
| Reproductive System Disease | 7.89E-03 |
| Gastrointestinal Disease | 8.75E-03 |
| Endocrine System Disorders | 7.89E-03 |
| c. Molecular and Cellular Functions |  |
| Cellular Assembly and Organization | 9.38E-03 |
| Cellular Function and Maintenance | 8.19E-03 |
| Cellular Movement | 6.13E-03 |
| Cellular Development | 9.38E-03 |
| Cellular Growth and Proliferation | 9.38E-03 |
| d. Physiological System Development and Function |  |
| Nervous System Development and Function | 9.09E-03 |
| Tissue Development | 9.38E-03 |
| Organismal Survival | 3.97E-04 |
| Organismal Development | 9.38E-03 |
| Embryonic Development | 9.38E-03 |

Based on network pathway analysis it was found that Neutrophil extracellular trap, Fc Receptor-mediated Phagocytosis in Macrophages and Monocytes and HGF signaling were found associated through dephosphorylation for regeneration of caudal fin tissue. Whereas mTOR signaling and Insulin signaling were associated with phosphorylation during regeneration (Figure 4a). Generation of tumour, frequency of tumour, incidence of tumour and carcinoma were found associated with dephosphorylation and infection of epithelial cells, embryonic cells and kidney cells association through phosphorylation are the most associated disease and functions (Figure 4b). Cancer, haematological & immunological disease and cancer, organismal injury & abnormalities were found to be the most associated canonical network pathways with the differentially expressed phosphoproteome (Figure 4c and 4d). A total 26 proteins such as ANO1, PRKCI, CCDC86, CLASP2, CSDE1, DDRGK1, DDX41, DOCK1, E2f, EIF4ENIF1, EML4, FAM76B, G3BP2, GTPase, HECW2, OSBPL8, PRKCI, PUM1, RANBP2, RCC2, SNW1, SRSF11, TCF, TFIP11, WBP4 and WDR7 were found undergoing differential phosphorylation and associated with cancer, haematological and immunological disease network pathway (Figure 4c). Similarly, for cancer, organismal injury and abnormalities canonical pathway a total of 26 proteins from the list were found to be associated and differentially phosphorylated (Figure 4d). The proteins include ACTB, AHNAK, CALD1, CCNY, DBN1, DSG2, EPB41L2, FAM83G, FKBP15, GBF1, KLC1, KLC2, LIMA1, MYH9, PAH, PP1 protein complex group, PLIN3, PPP1R12A, PPP4R3B RAPH1, SCIN, SLC4A1AP, SORT1, SSH3, SVIL, TRIM14 and VPS26B.

**Figure 4:**
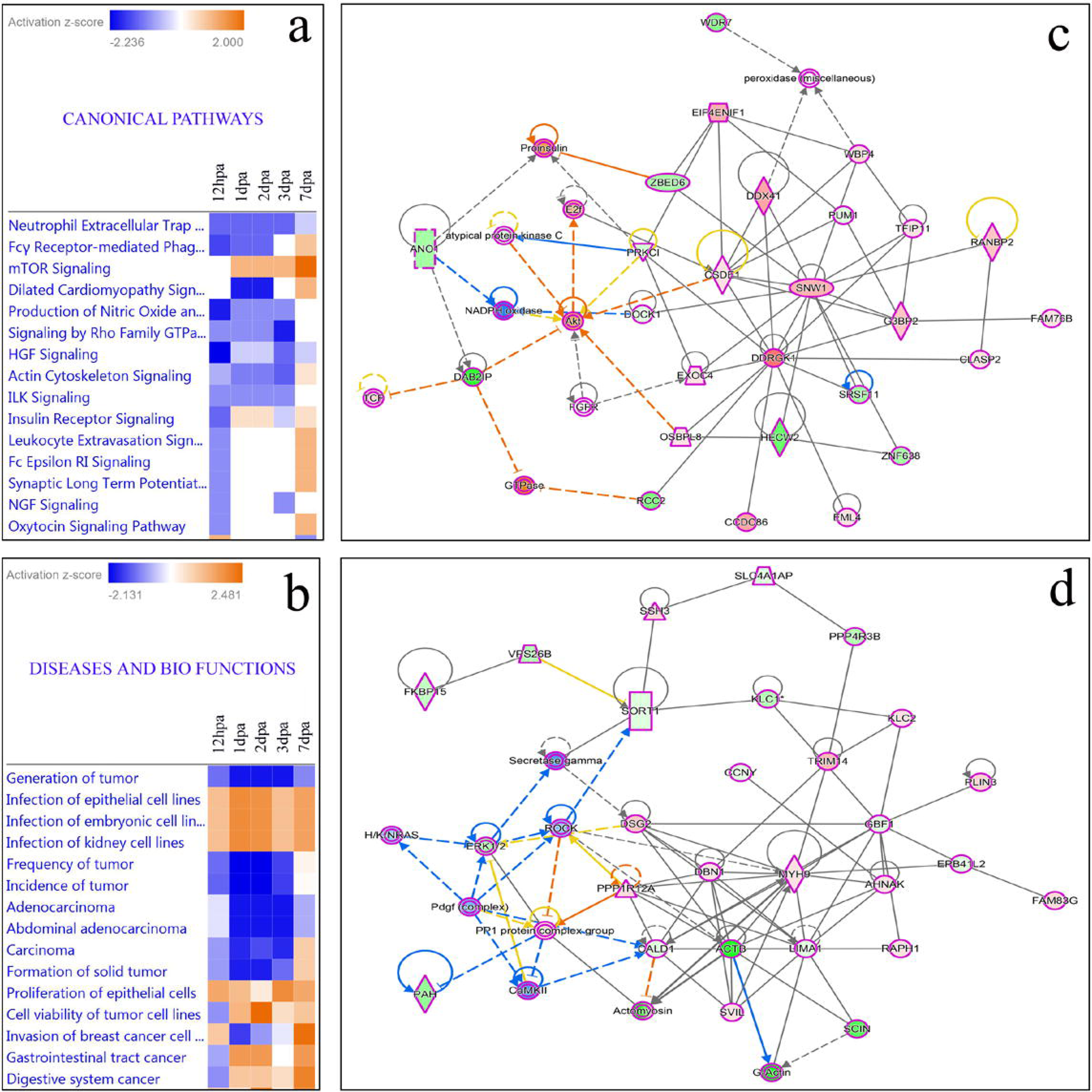
Canonical network pathways and Disease & Biofunctions pathways associated with the differentially expressed phosphoproteome for zebrafish caudal fin regeneration a. Heat map of top 15 canonical pathways. b. Heat map of disease and bio functions. c. Cancer, haematological & immunological disease canonical pathway d. Cancer, organismal injury & abnormalities canonical network pathways.

## Discussion

Phosphorylation, a pivotal regulatory post-translational modification, orchestrates cellular signaling and protein function. This study delves into the intricate landscape of phosphorylation events during zebrafish caudal fin tissue regeneration and the critical processes involved. Our phosphoproteomic analyses revealed dynamic changes in protein phosphorylation throughout the regenerative process. Notably, 74 proteins showed differential phosphorylation patterns via TiO2 column enrichment, while immunoprecipitation with phosphoserine, phosphothreonine and phosphotyrosine antibodies identified 440 proteins undergoing phosphorylation modification. The robust correlation (95%) between the two methods validates the accuracy of our findings.

### Phosphorylation and Regeneration

It is very interesting to observe the cascade of proteins undergoing phosphorylation and Dephosphorylation during the most orchestrated event of regeneration in zebrafish. Major proteins such as LIM domain and actin-binding protein 1-like, Exocyst complex component 7 isoform X10, Arsenite resistance protein 2, WW domain-binding protein 4, connector enhancer of kinase suppressor of ras 1 isoform X1, Ceruloplasmin precursor, serine/arginine-rich splicing factor 1A isoform X1 and cytoplasmic dynein 1 light intermediate chain 2 were found undergoing phosphorylation during the process of epimorphic regeneration in zebrafish.

LIM domain and actin-binding protein 1 (LIMA1) is involved in actin cytoskeleton regulation and dynamics. LIMA1 increases the number and size of actin stress fibers and inhibits membrane ruffling and also inhibits actin filament depolymerization, bundles actin filaments, delays filament nucleation and reduces formation of branched filaments^16^. An increased phosphorylation of LIMA1 was found to be associated throughout the regeneration process based on both TiO2 and antibody immunoprecipitation methods (figure 1). Cytoplasmic dynein 1 light intermediate chain 2 (dync1li2) was also actively phosphorylating during the process of regeneration. dync1li2 acts as one of several non-catalytic accessory components of the cytoplasmic dynein 2 complex (dynein-2 complex), a motor protein complex that drives the movement of cargos along microtubules within cilia and flagella in concert with the intraflagellar transport (IFT) system, facilitating the assembly of these organelles and also involved in the regulation of ciliary length^17^.

Phosphorylation of WW domain-binding protein 4 (WBP4) was found to be involved in pre-mRNA splicing as a component of the spliceosome, which also plays a role in cross-intron bridging of U1 and U2 snRNPs in the mammalian A complex^18^. Connector enhancer of kinase suppressor of ras 1 isoform X1 (cnksr1) involved in function as an adapter protein or regulator of Ras signaling pathways^19^ and serine/arginine-rich splicing factor 1A isoform X1 (srsf1a) involved in prevention of exon skipping, ensuring the accuracy of splicing and regulating alternative splicing were also undergoing phosphorylation during the regeneration process^20^. Ceruloplasmin belongs to the copper oxidase family and is mainly involved in copper transport. Additionally, it plays a role in ferrous and calcium ion transport and exhibits antioxidant activity. Ceruloplasmin is linked with ferroportin (encoded by SLC40A1) for iron transport. Impairment in the ceruloplasmin and ferroportin system is associated with NBIA (neurodegeneration with brain iron accumulation) and aceruloplasminemia, where ion-mediated oxidative stress is observed^21^. Our previous study demonstrated the predominant expression of other SLC family transport mechanisms in zebrafish caudal fin regeneration^5^.

### Dephosphorylation and Regeneration

As like phosphorylation, several proteins were observed undergoing dephosphorylation against the control which includes majorly disabled homolog 2-interacting protein (DAB2IP), H2B, PRDM2, MAP3K15, NDRG2 and annexin proteins such as ANXA1a, 2a, 11a and 5b. Disabled homolog 2-interacting protein (DAB2IP), a Ras GTPase-activating protein, is downregulated in several cancers and was dephosphorylated during the process of regeneration (Supplementary Table 2). DAB2IP modulates the balance between phosphatidylinositol 3-kinase (PI3K)-AKT-mediated cell survival and apoptosis stimulated kinase (MAP3K5)-JNK signaling pathways; sequesters both AKT1 and MAP3K5 counterbalances the activity of each kinase by modulating their phosphorylation status in response to pro-inflammatory stimuli^22^. DAB2IP mediates TNF-alpha-induced apoptosis activation by facilitating dissociation of inhibitor 14-3-3 from MAP3K5; recruits the PP2A phosphatase complex which dephosphorylates MAP3K5 on ‘Ser-966’, leading to the dissociation of 14-3-3 proteins and activation of the MAP3K5-JNK signaling pathway in endothelial cells.

The Core component of nucleosome histone H2B1/2 was found to be dephosphorylated during the process of regeneration. Nucleosomes wrap and compact DNA into chromatin, limiting DNA accessibility to the cellular machineries which require DNA as a template. Histones thereby play a central role in transcription regulation, DNA repair, DNA replication and chromosomal stability. DNA accessibility is regulated via a complex set of post-translational modifications of histones, also called as histone code, and nucleosome remodeling^23^. Protein NDRG2 contributes to the regulation of the Wnt signaling pathway. It down-regulates CTNNB1-mediated transcriptional activation of target genes, such as CCND1, and may thereby act as tumor suppressor^24^. Dephosphorylation of NDGR2 and NDRG4 were found most specifically during the structural regeneration stage of caudal fin (Supplementary Table 2).

It is well documented from our previous study that Annexin is highly upregulated during the process of regeneration^7–9^ and also undergoes dephosphorylation during the process of regeneration^7^. In this study it is further confirmed that ANXA1a, 2a, 5b and 11a undergoes dephosphorylation during the process of regeneration right from wound healing stage to structural regeneration stage impacting the strong association (Supplementary table 2).

### Conclusion

In conclusion, our investigation into the phosphorylation dynamics during zebrafish caudal fin regeneration has successfully identified crucial proteins, elucidated their dynamic phosphorylation states, and uncovered their pivotal roles in fundamental signaling pathways. This breakthrough significantly enhances our understanding of the intricate molecular mechanisms driving tissue regeneration. Importantly, the insight into these phosphorylation events not only augments our current knowledge but also establishes them as promising targets for advancing tissue regeneration.

## References

1. Gemberling, M., Bailey, T. J., Hyde, D. R. & Poss, K. D. The zebrafish as a model for complex tissue regeneration. Trends in Genetics vol. 29 611–620 (2013).

2. Akimenko, M. A., Marí-Beffa, M., Becerra, J. & Géraudie, J. Old questions, new tools, and some answers to the mystery of fin regeneration. Developmental Dynamics vol. 226 190–201 (2003).

3. Pfefferli, C. & Jaźwińska, A. The art of fin regeneration in zebrafish. Regeneration 2, 72– 83 (2015).

4. Sehring, I. M. & Weidinger, G. Recent advancements in understanding fin regeneration in zebrafish. Wiley Interdisciplinary Reviews: Developmental Biology vol. 9 (2020).

5. Banu, S. et al. Understanding the complexity of epimorphic regeneration in zebrafish caudal fin tissue: A transcriptomic and proteomic approach. Genomics 114, (2022).

6. Hou, Y., et al. Cellular diversity of the regenerating caudal fin. Sci. Adv 6(33):eaba2084 (2020).

7. Saxena, S. et al. Proteomic analysis of zebrafish caudal fin regeneration. Molecular and Cellular Proteomics 11, (2012).

8. Saxena, S. et al. Role of annexin gene and its regulation during zebrafish caudal fin regeneration. Wound Repair and Regeneration 24, 551–559 (2016).

9. Quoseena, M. et al. Functional role of annexins in zebrafish caudal fin regeneration – A gene knockdown approach in regenerating tissue. Biochimie 175, 125–131 (2020).

10. Jin, J. & Pawson, T. Modular evolution of phosphorylation-based signalling systems. Philosophical Transactions of the Royal Society B: Biological Sciences vol. 367 2540– 2555 (2012).

11. Higgins, L., Gerdes, H. & Cutillas, P. R. Principles of phosphoproteomics and applications in cancer research. Biochemical Journal vol. 480 403–420 (2023).

12. Zhang, M., Liu, C., Zhao, L., Zhang, X. & Su, Y. The Emerging Role of Protein Phosphatase in Regeneration. Life vol. 13 (2023).

13. Kwon, O. K. et al. Global analysis of phosphoproteome dynamics in embryonic development of zebrafish (Danio rerio). Proteomics 16, 136–149 (2016).

14. Hardman, G. et al. Strong anion exchangeLmediated phosphoproteomics reveals extensive human nonLcanonical phosphorylation. EMBO J 38, (2019).

15. Banu, S., Nagaraj, R. & Idris, M. M. A proteomic perspective and involvement of cytokines in SARS-CoV-2 infection. PLoS One 18, (2023).

16. Maul, R. S. et al. EPLIN regulates actin dynamics by cross-linking and stabilizing filaments. Journal of Cell Biology 160, 399–407 (2003).

17. Hamada, Y., Tsurumi, Y., Nozaki, S., Katoh, Y. & Nakayama, K. Interaction of WDR60 intermediate chain with TCTEX1D2 light chain of the dynein-2 complex is crucial for ciliary protein trafficking. Mol Biol Cell 29, 1628–1639 (2018).

18. Bertram, K. et al. Cryo-EM Structure of a Pre-catalytic Human Spliceosome Primed for Activation. Cell 170, 701–713.e11 (2017).

19. Therrien, M., Wong, A. M. & Rubin, G. M. *CNK*, a RAF-Binding Multidomain Protein Required for RAS Signaling Some Evi-Dence Indicates That RAF and MEK Are Part of a Complex Dominant-Negative Molecule When Separated from The. Cell vol. 95 (1998).

20. Zheng, X. et al. Serine/arginine-rich splicing factors: The bridge linking alternative splicing and cancer. International Journal of Biological Sciences vol. 16 2442–2453 Preprint at 10.7150/ijbs.46751 (2020).

21. Musci, G., Polticelli, F., & di Patti, M. C. B. Ceruloplasmin-ferroportin system of iron traffic in vertebrates. World journal of biological chemistry 5, 204 (2014).

22. Xie, D. et al. DAB2IP Coordinates Both PI3K-Akt and ASK1 Pathways for Cell Survival and Apoptosis. www.pnas.org/cgi/content/full/.

23. Luo, A. et al. H2B ubiquitination recruits FACT to maintain a stable altered nucleosome state for transcriptional activation. Nat Commun 14, (2023).

24. Hwang, J. et al. Crystal structure of the human N-Myc downstream-regulated gene 2 protein provides insight into its role as a tumor suppressor. Journal of Biological Chemistry 286, 12450–12460 (2011).

