## Supplemental Table 1 for "Exploration of phosphoproteomic association during epimorphic regeneration"

Supplementary Table 1: List of Proteins identified to differential phosphorylation based on TiO2 column enrichment assay.

| S.No | Accession | Description | Symbol | # PSMs | 12hpa | 1dpa | 2dpa | 3dpa | 7dpa |
| --- | --- | --- | --- | --- | --- | --- | --- | --- | --- |
| 1 | NP_861425.1 | annexin A1c | ANXA1c | 2 | -1.8 | -2.2 | -1.9 | -3.1 | -0.5 |
| 2 | XP_005165804.1 | disabled homolog 2-interacting protein | DAB2 | 8 | -1.4 | -1.3 | -1.0 | -0.8 | -1.8 |
| 3 | NP_861426.1 | annexin A2a | ANXA2a | 9 | -1.2 | -1.7 | -0.9 | -1.8 | -0.9 |
| 4 | XP_021333836.1 | 4-trimethylaminobutyraldehyde dehydrogenase isoform X1 | ALDH9A1 | 2 | -1.2 | -2.0 | -1.6 | -2.3 | -1.2 |
| 5 | XP_021334590.1 | uncharacterized protein col6a3 isoform X1 | COL6A3 | 13 | -1.2 | -1.0 | -1.5 | -1.6 | -2.2 |
| 6 | NP_835232.2 | scinderin like a | scinla | 3 | -1.1 | -2.2 | -1.4 | -3.2 | -0.9 |
| 7 | NP_001070922.1 | keratin 94 | KRT94 | 43 | -1.0 | 0.0 | 3.4 | 2.8 | 2.4 |
| 8 | XP_021333143.1 | calcium/calmodulin-dependent protein kinase type II delta 1 chain isoform X6 | camk2d1 | 10 | -1.0 | -1.8 | -1.6 | -2.5 | -1.8 |
| 9 | NP_861422.2 | annexin A5b | ANXA5b | 2 | -0.9 | -1.5 | -1.8 | -2.2 | -1.6 |
| 10 | NP_001121753.1 | dual specificity mitogen-activated protein kinase kinase 2b | MAP2K2B | 7 | -0.9 | -1.1 | -1.1 | -1.3 | -1.5 |
| 11 | NP_001070115.2 | XIAP-associated factor 1 | XAF1 | 3 | -0.9 | -1.5 | -1.6 | -2.4 | -1.8 |
| 12 | QFX67701.1 | transforming growth factor beta-activated kinase 1b | TAK1 | 8 | -0.8 | -0.9 | -1.1 | -1.0 | -0.7 |
| 13 | NP_956845.1 | phenylalanine-4-hydroxylase | PAH | 22 | -0.8 | -0.4 | -1.9 | -1.9 | -1.5 |
| 14 | XP_009293997.1 | protein NLRC3 | NLRC3 | 5 | -0.6 | -0.9 | -1.1 | -0.5 | -0.7 |
| 15 | NP_878308.2 | mitogen-activated protein kinase 1 | MAPK1 | 5 | -0.6 | -0.6 | -0.7 | -1.1 | 0.4 |
| 16 | XP_009303246.1 | uncharacterized protein LOC445086 isoform X1 | zgc:92380 | 24 | -0.5 | -1.1 | -1.1 | -1.6 | -1.7 |
| 17 | XP_005157568.1 | vacuolar protein sorting-associated protein 26B isoform X1 | VPS26B | 4 | -0.5 | -0.3 | -2.9 | -4.2 | -2.7 |
| 18 | Q90XF2.2 | Protein kinase C iota type | PRKCI | 7 | -0.3 | 1.2 | 2.2 | 1.2 | 2.5 |
| 19 | XP_005172514.1 | uncharacterized protein LOC393431 isoform X1 | LOC393431 | 24 | -0.2 | -1.0 | -1.9 | -2.3 | -1.6 |
| 20 | NP_001243104.1 | E3 ubiquitin-protein ligase BRE1A | RNF20 | 7 | -0.2 | 0.8 | 1.0 | 1.2 | 0.8 |
| 21 | XP_021335365.1 | LIM domain and actin-binding protein 1-like | LIMA1 | 34 | -0.1 | 0.9 | 1.1 | 0.9 | 1.3 |
| 22 | NP_001017624.1 | 26S proteasome non-ATPase regulatory subunit 4b | psmd4b | 15 | -0.1 | -0.7 | -1.0 | -0.9 | -1.0 |
| 23 | XP_005167369.1 | AP2-associated protein kinase 1 isoform X1 | AAK1 | 2 | 0.0 | -0.1 | 0.6 | 1.0 | 1.1 |
| 24 | NP_571077.2 | lamin-B2 | LMNB2 | 10 | 0.0 | 0.5 | 1.3 | 1.3 | 0.5 |
| 25 | NP_001001944.2 | 5'-3' exoribonuclease 2 | XRN2 | 6 | 0.0 | 1.0 | 1.3 | 1.4 | 2.6 |
| 26 | NP_001002201.1 | drebrin-like a | DBNL | 6 | 0.0 | -2.0 | -1.4 | -0.2 | 0.0 |
| 27 | XP_009293975.2 | NACHT, LRR and PYD domains-containing protein 3-like | NLRP3 | 4 | 0.1 | -2.2 | -2.0 | -0.9 | -0.4 |
| 28 | NP_997819.1 | tyrosine-protein phosphatase non-receptor type 2 | ptpn2b | 6 | 0.1 | 1.3 | 0.7 | 0.9 | 2.2 |
| 29 | XP_009304816.1 | supervillin isoform X1 | SVIL | 8 | 0.2 | 1.0 | 1.5 | 1.5 | 1.2 |
| 30 | NP_997764.1 | interleukin enhancer-binding factor 3 homolog | ilf3b | 3 | 0.2 | -0.1 | 1.2 | 0.8 | 0.7 |
| 31 | XP_021336562.1 | uncharacterized protein LOC563946 isoform X1 | zgc:136930 | 49 | 0.2 | -0.5 | -1.2 | -2.3 | -1.7 |
| 32 | NP_001070846.1 | nuclear-interacting partner of ALK | ZC3HC1 | 29 | 0.3 | 0.9 | 0.4 | 1.1 | 1.2 |
| 33 | NP_998633.1 | aquaporin-3a | AQP3A | 9 | 0.3 | -0.2 | -0.7 | -1.4 | -2.1 |
| 34 | XP_005165358.1 | Golgi phosphoprotein 3 isoform X1 | golph3a | 12 | 0.3 | 0.9 | 1.2 | 1.7 | 1.2 |
| 35 | NP_001074055.1 | protein Niban 2a | NIBAN2A | 11 | 0.3 | 1.0 | 0.8 | 1.0 | 1.3 |
| 36 | NP_001002721.1 | tuftelin-interacting protein 11 | TFIP11 | 6 | 0.4 | 1.6 | 2.1 | 2.8 | 0.7 |
| 37 | AAM34668.1 | arsenite resistance protein 2 / Serrate RNA effector molecule homolog | SRRT | 10 | 0.4 | 1.1 | 1.2 | 1.1 | 0.6 |
| 38 | XP_005173034.1 | exocyst complex component 7 isoform X10 | EXOC7 | 7 | 0.4 | 1.3 | 0.9 | 1.2 | 1.0 |
| 39 | XP_003200294.1 | SWI/SNF complex subunit SMARCC1 isoform X1 | SMARCC1 | 43 | 0.4 | 2.2 | 1.0 | 1.7 | 1.0 |
| 40 | B5DE31.1 | Y-box-binding protein 1 | YBX1 | 99 | 0.4 | 0.7 | 1.7 | 1.2 | 1.3 |
| 41 | XP_005163413.1 | phospholipase C, delta 1b isoform X1 | plcd1b | 6 | 0.5 | 1.1 | 2.0 | 1.5 | 2.7 |
| 42 | NP_001139027.1 | ribosome-binding protein 1a | RRBP1A | 10 | 0.5 | 0.7 | 0.5 | 2.6 | 2.8 |
| 43 | XP_009291361.1 | connector enhancer of kinase suppressor of ras 1 isoform X1 | CNKSR1 | 3 | 0.5 | 0.9 | 1.4 | 1.5 | 0.8 |
| 44 | XP_021333866.1 | cold shock domain-containing protein E1 isoform X1 | CSDE1 | 22 | 0.5 | 0.9 | 1.1 | 1.1 | 0.6 |
| 45 | NP_001018530.1 | WW domain-binding protein 4 | WBP4 | 4 | 0.6 | 1.6 | 1.1 | 1.0 | 1.5 |
| 46 | XP_021334370.1 | CLIP-associating protein 1 isoform X1 | CLASP1 | 3 | 0.6 | 1.4 | 1.0 | 1.8 | 0.9 |
| 47 | XP_005157472.1 | serine/arginine-rich splicing factor 1A isoform X1 | SRSF1A | 15 | 0.6 | 1.8 | 1.2 | 2.0 | -1.6 |
| 48 | XP_005166071.1 | eukaryotic translation initiation factor 4B isoform X1 | eif4bb | 33 | 0.6 | 1.0 | 1.0 | 0.9 | 0.1 |
| 49 | XP_021330670.1 | protein phosphatase 1 regulatory subunit 12A isoform X1 | PPP1R12A | 9 | 0.6 | 1.1 | 1.3 | 1.4 | 0.8 |
| 50 | NP_001002749.1 | tyrosinase-related protein 1b precursor | TYRP1B | 5 | 0.7 | 0.2 | 2.3 | 1.5 | 3.7 |
| 51 | NP_001103951.1 | perilipin 6 | PLIN3 | 4 | 0.7 | 0.2 | 0.1 | -1.4 | -1.4 |
| 52 | XP_009301712.1 | transcription factor 12 isoform X4 | TCF12 | 2 | 0.7 | 0.1 | 1.2 | 2.0 | 1.5 |
| 53 | NP_001092707.2 | eukaryotic translation initiation factor 4Ba | EIF4BA | 5 | 0.8 | 0.9 | 0.8 | 1.0 | 1.2 |
| 54 | XP_021334470.1 | E3 SUMO-protein ligase RanBP2 isoform X1 | RANBP2 | 9 | 0.9 | 2.4 | 0.8 | 2.4 | 4.7 |
| 55 | XP_021325628.1 | eukaryotic translation initiation factor 4 gamma 3 isoform X1 | EIF4G3 | 2 | 1.0 | 0.7 | 1.2 | 1.3 | 1.9 |
| 56 | NP_571655.2 | 60S acidic ribosomal protein P0 | RPLP0 | 19 | 1.0 | 1.5 | 1.0 | 2.4 | 1.6 |
| 57 | XP_001338671.2 | caldesmon, smooth muscle-like isoform X1 | CALD1 | 24 | 1.0 | 1.7 | 2.2 | 2.0 | -1.6 |
| 58 | XP_005159421.1 | ets2 repressor factor isoform X1 | erf | 3 | 1.1 | 0.8 | 1.5 | 1.2 | 1.2 |
| 59 | NP_956279.1 | coiled-coil domain-containing protein 6a | CCDC6A | 4 | 1.2 | 1.7 | 1.2 | 1.4 | 1.2 |
| 60 | NP_001002864.1 | SNW domain-containing protein 1 | SNW1 | 14 | 1.2 | 1.7 | 1.6 | 2.0 | 2.0 |
| 61 | NP_997732.2 | DNA replication licensing factor MCM3 | MCM3 | 4 | 1.3 | 0.8 | 2.1 | 2.1 | 1.3 |
| 62 | XP_021332540.1 | eukaryotic translation initiation factor 4E transporter isoform X1 | EIF4ENIF1 | 6 | 1.3 | 2.0 | 1.9 | 2.0 | 2.9 |
| 63 | XP_021326591.1 | vitellogenin-like | VTG | 38 | 1.3 | 2.0 | 1.4 | 4.0 | 0.0 |
| 64 | NP_001138831.1 | coiled-coil domain-containing protein 86 | CCDC86 | 7 | 1.4 | 1.4 | 2.8 | 1.4 | 1.5 |
| 65 | XP_005164248.1 | calponin homology domain-containing protein DDB_G0272472 isoform X1 | LRCH1 | 6 | 1.5 | 1.7 | 2.1 | 1.6 | 1.2 |
| 66 | NP_997970.1 | smoothelin, like | SMTN | 9 | 1.7 | 1.1 | 2.4 | 0.8 | 1.4 |
| 67 | NP_001268717.1 | A-kinase anchor protein 12b isoform 1 | AKAP12B | 8 | 1.8 | 4.0 | 2.7 | 3.6 | 5.2 |
| 68 | XP_005170607.1 | kininogen-1 isoform X1 | KNG1 | 102 | 1.8 | 2.8 | 2.8 | 2.4 | 2.8 |
| 69 | NP_001038745.1 | plakophilin-3a | PKP3A | 54 | 2.0 | 2.2 | 2.1 | 2.1 | 2.0 |
| 70 | XP_009301350.1 | splicing factor 1 isoform X1 Branchpoint-bridging protein | sf1 | 10 | 2.1 | 2.7 | 3.4 | 4.0 | 3.8 |
| 71 | CAP09627.1 | novel protein | fxyd3 | 10 | 2.1 | 1.3 | 1.8 | 2.5 | 1.3 |
| 72 | NP_001038362.3 | vitellogenin 1 precursor | VTG1 | 40 | 2.2 | 3.1 | 1.5 | 4.7 | -0.7 |
| 73 | NP_956587.2 | DDRGK domain-containing protein 1 precursor | DDRGK1 | 2 | 2.2 | 3.5 | 3.0 | 3.8 | 3.3 |
| 74 | XP_021324336.1 | uridine-cytidine kinase-like 1 isoform X1 | UCKL1 | 4 | 3.2 | 4.0 | 4.3 | 4.2 | 1.9 |
