## Supplemental Table 2 for "Exploration of phosphoproteomic association during epimorphic regeneration"

Supplementary Table 2: List of Proteins identified to differential phosphorylation based on immunoprecipitation assay involving pSerine, pThreonine and pTyrosine antibodies.

| S. No | **Accession** | **Description** | **Symbol** | **Type of Phosphorylation** | **Identified from TiO2** | **Mean Expression** | | | | | **SEM** | | | | |
| --- | --- | --- | --- | --- | --- | --- | --- | --- | --- | --- | --- | --- | --- | --- | --- |
|  |  |  |  |  |  | **12hpa** | **1dpa** | **2dpa** | **3dpa** | **7dpa** | **12hpa** | **1dpa** | **2dpa** | **3dpa** | **7dpa** |
| 1 | XP_005167369.1 | AP2-associated protein kinase 1 isoform X1 | AAK1 | Serine | YES | 0.0 | -0.1 | 0.6 | 0.9 | 1.0 | NA | NA | NA | NA | NA |
| 2 | XP_009295502.1 | abl interactor 1 isoform X1 | abi1a | Threonine | No | 0.5 | 0.1 | -0.5 | 0.7 | 2.1 | NA | NA | NA | NA | NA |
| 3 | XP_009304888.1 | actin-binding LIM protein 1 isoform X1 | ablim1b | Serine, Threonine | No | 0.1 | 0.2 | 0.0 | -0.2 | -0.8 | 0.0 | 0.3 | 0.4 | 0.2 | 0.5 |
| 4 | XP_009305002.1 | actin-binding LIM protein 2 isoform X1 | ABLIM2 | Serine | No | 0.0 | -0.1 | -0.2 | -0.4 | -0.6 | NA | NA | NA | NA | NA |
| 5 | XP_021324516.1 | alpha-adducin isoform X1 | add1 | Serine | No | 0.6 | 1.2 | 0.7 | 0.4 | 0.9 | NA | NA | NA | NA | NA |
| 6 | XP_017208635.1 | gamma-adducin isoform X1 | add3a | Serine | No | 0.3 | 1.8 | 1.0 | 0.7 | 0.6 | NA | NA | NA | NA | NA |
| 7 | XP_021324351.1 | afadin isoform X1 | AFDN | Serine, Threonine | No | 0.7 | 1.8 | 0.6 | 1.5 | 0.8 | 0.5 | 0.1 | 0.2 | 0.1 | 0.9 |
| 8 | XP_009300560.1 | arf-GAP with GTPase, ANK repeat and PH domain-containing protein 1 isoform X1 | AGAP1 | Serine | No | 0.6 | -0.4 | 0.3 | 0.7 | 0.4 | NA | NA | NA | NA | NA |
| 9 | XP_009295860.1 | arf-GAP with GTPase, ANK repeat and PH domain-containing protein 3 isoform X1 | AGAP3 | Serine | No | 0.9 | -0.2 | 0.7 | 0.4 | 0.5 | NA | NA | NA | NA | NA |
| 10 | NP_998384.2 | A-kinase anchor protein 8-like | akap8l | Threonine | No | 1.2 | 2.1 | 0.0 | 1.8 | 0.2 | NA | NA | NA | NA | NA |
| 11 | XP_021333836.1 | 4-trimethylaminobutyraldehyde dehydrogenase isoform X1 | aldh9a1a.2 | Serine | YES | -0.6 | -2.0 | -1.1 | -2.3 | -1.4 | NA | NA | NA | NA | NA |
| 12 | NP_998380.1 | aldolase a, fructose-bisphosphate, b | aldoab | Threonine | No | -0.1 | 0.2 | 0.6 | -0.6 | -0.3 | NA | NA | NA | NA | NA |
| 13 | NP_956142.1 | AMP deaminase 3b | ampd3b | Serine | No | -0.2 | -0.2 | -0.7 | -0.8 | -0.8 | NA | NA | NA | NA | NA |
| 14 | XP_009290458.1 | ankyrin repeat and IBR domain-containing protein 1 | ANKIB1 | Serine | No | 0.1 | 0.9 | 0.9 | 1.4 | 0.3 | NA | NA | NA | NA | NA |
| 15 | XP_009301746.1 | anoctamin-1 isoform X1 | ano1a | Serine | No | -0.6 | -0.6 | -1.2 | -1.6 | -1.4 | NA | NA | NA | NA | NA |
| 16 | NP_861430.2 | annexin A11a isoform 1 | anxa11a | Serine, Threonine, Tyrosine | No | -0.7 | -0.6 | -0.1 | -1.2 | 0.2 | 0.1 | 0.2 | 0.1 | 0.3 | 0.1 |
| 17 | NP_861423.1 | annexin A1a | anxa1a | Serine, Threonine, Tyrosine | No | -0.6 | 0.4 | -0.2 | -1.2 | 0.2 | 0.1 | 0.2 | 0.1 | 0.3 | 0.1 |
| 18 | NP_861426.1 | annexin A2a | anxa2a | Serine, Threonine, Tyrosine | YES | -1.1 | -2.0 | -1.1 | -1.8 | -0.7 | 0.1 | 0.6 | 0.6 | 0.7 | 0.2 |
| 19 | NP_861422.2 | annexin A5b | anxa5b | Threonine, Tyrosine | YES | -0.7 | -1.1 | -1.5 | -1.6 | -1.4 | 0.2 | 0.2 | 0.1 | 0.3 | 0.1 |
| 20 | NP_001093614.2 | apolipoprotein A-Ib precursor | apoa1b | Serine, Threonine, Tyrosine | No | 0.0 | -0.6 | -0.2 | -1.3 | -1.2 | 0.1 | 0.2 | 0.1 | 0.3 | 0.1 |
| 21 | NP_998633.1 | aquaporin-3a | aqp3a | Serine | YES | 0.2 | 0.1 | -0.4 | -1.1 | -1.1 | NA | NA | NA | NA | NA |
| 22 | NP_958867.1 | archain 1a | arcn1a | Threonine | No | 0.5 | 2.2 | 2.1 | 2.2 | 3.3 | NA | NA | NA | NA | NA |
| 23 | XP_005159305.1 | ADP-ribosylation factor GTPase-activating protein 2 isoform X1 | arfgap2 | Serine | No | 0.1 | -1.0 | 1.0 | 0.0 | 1.1 | NA | NA | NA | NA | NA |
| 24 | NP_001032507.1 | ADP-ribosylation factor GTPase-activating protein 2 | arfgap2 | Threonine | No | 0.4 | 0.5 | 2.0 | 0.8 | 1.9 | NA | NA | NA | NA | NA |
| 25 | XP_005170948.1 | arfaptin-1 isoform X1 | arfip1 | Serine | No | -0.7 | 0.2 | 0.4 | 0.1 | -0.4 | NA | NA | NA | NA | NA |
| 26 | XP_009297597.1 | rho GTPase-activating protein 17 isoform X1 | arhgap17b | Serine | No | -2.4 | 2.9 | 0.2 | 1.0 | 2.1 | NA | NA | NA | NA | NA |
| 27 | XP_001918968.3 | rho GTPase-activating protein 21 isoform X1 | ARHGAP21 | Tyronsine | No | 0.4 | 0.4 | 1.2 | 0.4 | 0.8 | NA | NA | NA | NA | NA |
| 28 | XP_001343636.1 | rho GTPase-activating protein 35 | ARHGAP35 | Serine, Threonine | No | -0.3 | -0.7 | -0.7 | -0.7 | -0.6 | 0.0 | 0.1 | 0.1 | 0.2 | 0.1 |
| 29 | XP_001919378.4 | rho GTPase-activating protein 45-like isoform X1 | ARHGAP45 | Threonine | No | 0.2 | 1.6 | 0.3 | 1.4 | 0.8 | NA | NA | NA | NA | NA |
| 30 | XP_001336016.1 | rho GTPase-activating protein 5 | ARHGAP5 | Serine | No | -0.3 | 0.4 | 0.5 | 1.1 | 0.4 | NA | NA | NA | NA | NA |
| 31 | XP_017210203.1 | rho guanine nucleotide exchange factor 7 isoform X1 | arhgef7b | Serine | No | -0.4 | -0.7 | -0.8 | -1.5 | -0.2 | NA | NA | NA | NA | NA |
| 32 | XP_021323975.1 | AT-rich interactive domain-containing protein 1A | ARID1A | Serine, Threonine | No | -0.8 | -0.2 | -0.7 | -0.4 | -0.2 | 0.0 | 0.1 | 0.1 | 0.2 | 0.1 |
| 33 | NP_001121821.1 | ataxin-2 | atxn2 | Threonine | No | 0.9 | 1.4 | 1.5 | 1.3 | 0.6 | NA | NA | NA | NA | NA |
| 34 | XP_005163856.1 | ataxin-2-like protein isoform X1 | atxn2l | Serine, Threonine | No | 0.1 | -0.1 | 0.3 | 0.4 | 0.4 | 0.2 | 0.3 | 0.1 | 0.1 | 0.0 |
| 35 | XP_005163812.1 | brain-specific angiogenesis inhibitor 1-associated protein 2 isoform X1 | baiap2a | Serine | No | 0.2 | 0.0 | -0.3 | -0.6 | -0.2 | NA | NA | NA | NA | NA |
| 36 | XP_009304868.1 | BAI1-associated protein 2-like 1 isoform X1 | baiap2l1a | Serine, Threonine | No | -0.1 | -0.8 | -0.5 | -0.6 | -0.3 | 0.1 | 0.2 | 0.1 | 0.2 | 0.1 |
| 37 | XP_021323361.1 | breast cancer anti-estrogen resistance protein 1 isoform X1 | bcar1 | Serine, Threonine | No | -0.3 | -0.1 | -0.2 | -0.3 | 0.5 | 0.1 | 0.2 | 0.2 | 0.3 | 0.1 |
| 38 | NP_998225.2 | [3-methyl-2-oxobutanoate dehydrogenase [lipoamide]] kinase, mitochondrial | bckdk | Serine, Threonine, Tyrosine | No | -0.3 | -0.7 | 0.4 | -1.0 | -0.3 | 0.1 | 0.2 | 0.1 | 0.3 | 0.1 |
| 39 | NP_001116861.1 | bromodomain-containing protein 3a | brd3a | Serine, Threonine | No | -0.9 | 1.2 | -0.4 | 1.6 | -1.2 | 0.2 | 0.1 | 0.0 | 0.1 | 0.0 |
| 40 | NP_001005945.1 | telomere length and silencing protein 1 homolog | C5H9orf78 | Serine, Threonine | No | 0.6 | 1.3 | 1.2 | 1.4 | 0.7 | 0.0 | 0.2 | 0.2 | 0.2 | 0.1 |
| 41 | XP_021333143.1 | calcium/calmodulin-dependent protein kinase type II delta 1 chain isoform X6 | camk2d1 | Serine, Threonine | YES | -1.0 | -1.2 | -1.1 | -1.9 | -1.7 | 0.1 | 0.3 | 0.3 | 0.3 | 0.1 |
| 42 | XP_021331879.1 | calmodulin-regulated spectrin-associated protein 1-B isoform X1 | camsap1b | Threonine | No | 2.4 | 3.9 | 0.2 | 2.9 | 2.4 | NA | NA | NA | NA | NA |
| 43 | NP_998613.1 | calnexin precursor | canx | Serine | No | 0.1 | 1.4 | 1.8 | 1.1 | 2.2 | NA | NA | NA | NA | NA |
| 44 | NP_001108021.1 | caveolae-associated protein 1b | cavin1b | Serine, Threonine | No | -0.1 | 0.3 | -0.2 | -0.1 | -1.8 | 0.0 | 0.1 | 0.3 | 0.0 | 0.1 |
| 45 | NP_001263999.1 | caveolae-associated protein 2a | cavin2a | Serine | No | -0.9 | -0.7 | -1.2 | -1.1 | -1.1 | NA | NA | NA | NA | NA |
| 46 | NP_956279.1 | coiled-coil domain-containing protein 6a | ccdc6a | Serine | YES | 1.2 | 1.7 | 1.2 | 1.4 | 1.1 | NA | NA | NA | NA | NA |
| 47 | NP_001138831.1 | coiled-coil domain-containing protein 86 | ccdc86 | Serine | YES | 0.6 | 1.4 | 2.8 | 1.4 | 1.4 | NA | NA | NA | NA | NA |
| 48 | XP_005157707.1 | coiled-coil domain-containing protein 9 isoform X1 | ccdc9 | Serine, Threonine | No | -0.5 | -0.2 | 0.3 | 0.0 | 0.0 | 0.0 | 0.1 | 0.1 | 0.2 | 0.1 |
| 49 | NP_001002315.1 | cerebral cavernous malformations protein 2 homolog | ccm2 | Serine | No | 0.1 | -0.1 | 0.5 | -0.2 | 0.1 | NA | NA | NA | NA | NA |
| 50 | XP_009295540.1 | cyclin-Y isoform X1 | CCNY | Serine | No | -0.2 | -0.6 | -0.2 | -0.3 | -0.5 | NA | NA | NA | NA | NA |
| 51 | NP_001008583.2 | CD2-associated protein | cd2ap | Serine, Threonine | No | -0.1 | -0.6 | -0.1 | -0.1 | 0.0 | 0.2 | 0.3 | 0.3 | 0.0 | 0.0 |
| 52 | NP_957255.1 | CD2 antigen cytoplasmic tail-binding protein 2 | cd2bp2 | Serine | No | 0.1 | 0.5 | 2.1 | 0.1 | 1.9 | NA | NA | NA | NA | NA |
| 53 | XP_021324168.1 | serine/threonine-protein kinase MRCK beta isoform X2 | cdc42bpb | Serine, Threonine | No | -0.4 | -0.2 | -0.1 | -0.5 | -0.1 | 0.1 | 0.2 | 0.2 | 0.3 | 0.1 |
| 54 | XP_003199689.1 | cdc42 effector protein 4 | CDC42EP4 | Serine | No | 0.6 | 0.2 | 0.6 | -0.3 | 0.1 | NA | NA | NA | NA | NA |
| 55 | NP_997729.1 | cyclin-dependent kinase 1 | cdk1 | Threonine, Tyrosine | No | 0.4 | 2.0 | 2.7 | 2.1 | 0.6 | 0.2 | 0.2 | 0.1 | 0.3 | 0.1 |
| 56 | NP_957480.1 | phosphatidate cytidylyltransferase 2 | cds2 | Serine, Threonine | No | -0.1 | 0.1 | 0.2 | -0.8 | -0.8 | 0.0 | 0.1 | 0.1 | 0.2 | 0.1 |
| 57 | XP_021324631.1 | collagen type IV alpha-3-binding protein isoform X1 | CERT1 | Serine | No | 0.5 | -0.9 | -1.0 | 0.8 | 0.3 | NA | NA | NA | NA | NA |
| 58 | NP_001186119.1 | complement factor H precursor | cfh | Threonine | No | -0.2 | -0.5 | -0.7 | -0.1 | 0.1 | NA | NA | NA | NA | NA |
| 59 | NP_998069.1 | charged multivesicular body protein 2b | chmp2bb | Serine | No | 0.0 | 0.0 | 0.5 | -0.8 | -0.5 | NA | NA | NA | NA | NA |
| 60 | XP_021334370.1 | CLIP-associating protein 1 isoform X1 | clasp1a | Serine | YES | 0.6 | 1.4 | 1.0 | 1.8 | 0.8 | NA | NA | NA | NA | NA |
| 61 | NP_001315188.1 | CLIP-associating protein 2 isoform 1 | clasp2 | Serine | No | 0.5 | 2.7 | 2.3 | 3.0 | -0.5 | NA | NA | NA | NA | NA |
| 62 | XP_009300267.1 | CAP-Gly domain-containing linker protein 1 isoform X1 | CLIP1 | Serine, Threonine | No | -1.0 | -0.3 | -1.8 | -0.9 | -1.7 | 0.0 | 0.0 | 0.2 | 0.1 | 0.1 |
| 63 | XP_005157480.1 | clathrin, heavy chain b (Hc) isoform X1 | cltcb | Serine, Threonine, Tyrosine | No | -0.3 | -0.3 | -0.3 | -0.2 | -0.5 | 0.1 | 0.2 | 0.1 | 0.3 | 0.1 |
| 64 | XP_009291361.1 | connector enhancer of kinase suppressor of ras 1 isoform X1 | cnksr1 | Serine, Threonine | YES | 0.9 | 1.0 | 1.6 | 1.7 | 0.8 | 0.3 | 0.1 | 0.1 | 0.2 | 0.1 |
| 65 | XP_005160168.1 | collagen, type XVII, alpha 1a isoform X1 | col17a1a | Threonine | No | -0.8 | -2.4 | -3.7 | -1.8 | -0.2 | NA | NA | NA | NA | NA |
| 66 | NP_892013.2 | collagen alpha-2(I) chain precursor | col1a2 | Serine, Threonine, Tyrosine | No | -1.5 | -1.9 | -2.2 | -2.0 | -1.3 | 0.2 | 0.2 | 0.1 | 0.3 | 0.1 |
| 67 | XP_021335547.1 | collagen alpha-2(VI) chain isoform X1 | COL6A2 | Threonine, Tyrosine | No | -0.6 | -1.2 | -0.6 | -1.2 | -0.3 | 0.2 | 0.2 | 0.1 | 0.3 | 0.1 |
| 68 | NP_571877.1 | ceruloplasmin precursor | cp | Serine, Threonine, Tyrosine | No | 0.1 | 0.6 | 1.2 | 0.2 | 0.9 | 0.2 | 0.2 | 0.1 | 0.3 | 0.1 |
| 69 | XP_021333866.1 | cold shock domain-containing protein E1 isoform X1 | csde1 | Serine, Threonine | YES | 0.6 | 1.0 | 1.2 | 1.3 | 0.6 | 0.0 | 0.2 | 0.1 | 0.2 | 0.1 |
| 70 | NP_571531.1 | catenin alpha-1 | ctnna1 | Serine, Threonine | No | 0.0 | -0.3 | 0.0 | -0.4 | 0.3 | 0.0 | 0.2 | 0.1 | 0.2 | 0.1 |
| 71 | XP_005157888.1 | catenin beta-1 isoform X1 | ctnnb1 | Serine, Threonine | No | -0.3 | -0.3 | 0.9 | -1.8 | 1.0 | 0.0 | 0.1 | 0.1 | 0.2 | 0.1 |
| 72 | XP_009291693.1 | catenin delta-1 isoform X3 | CTNND1 | Serine | No | 0.9 | 0.1 | 0.4 | -0.1 | 0.4 | NA | NA | NA | NA | NA |
| 73 | XP_005167394.1 | drebrin-like protein isoform X1 | dbnlb | Serine | No | -0.3 | -1.1 | 0.6 | -1.1 | 0.9 | NA | NA | NA | NA | NA |
| 74 | NP_956587.2 | DDRGK domain-containing protein 1 precursor | ddrgk1 | Serine | YES | 2.3 | 3.5 | 3.0 | 3.7 | 3.2 | NA | NA | NA | NA | NA |
| 75 | XP_005166610.1 | DENN domain-containing protein 4C isoform X1 | DENND4C | Serine | No | -0.2 | -0.2 | -0.2 | 0.2 | 0.4 | NA | NA | NA | NA | NA |
| 76 | NP_955917.1 | DnaJ (Hsp40) homolog, subfamily C, member 5 gamma a | dnajc5ga | Serine, Threonine | No | -0.5 | -0.1 | 0.8 | -0.4 | 0.7 | 0.0 | 0.1 | 0.1 | 0.2 | 0.1 |
| 77 | NP_001103481.1 | dedicator of cytokinesis protein 1 | dock1 | Serine, Threonine | No | -0.2 | 0.0 | -0.4 | -0.1 | -0.8 | 0.0 | 0.1 | 0.1 | 0.2 | 0.1 |
| 78 | NP_001018348.1 | dihydropyrimidinase-related protein 3 | dpysl3 | Serine, Threonine | No | -0.2 | -0.1 | -0.5 | -0.5 | -0.3 | 0.0 | 0.3 | 0.1 | 0.4 | 0.1 |
| 79 | XP_001919901.3 | desmoplakin isoform X1 | DSP | Serine, Threonine, Tyrosine | No | 0.5 | -0.5 | -0.1 | 0.3 | 0.5 | 0.6 | 0.4 | 0.2 | 0.7 | 0.4 |
| 80 | XP_021324396.1 | desmoplakin isoform X1 | DSP | Serine, Tyrosine | No | -0.3 | -0.2 | -0.3 | -0.4 | -0.5 | 0.1 | 0.1 | 0.1 | 0.3 | 0.1 |
| 81 | NP_001017669.2 | cytoplasmic dynein 1 light intermediate chain 2 | dync1li2 | Serine, Threonine, Tyrosine | No | 1.2 | 1.1 | 1.3 | 1.5 | 0.7 | 0.1 | 0.2 | 0.1 | 0.3 | 0.1 |
| 82 | XP_005158271.1 | dual specificity tyrosine-phosphorylation-regulated kinase 1B isoform X1 | dyrk1b | Tyronsine | No | 0.0 | 0.3 | 0.2 | 0.9 | -0.3 | NA | NA | NA | NA | NA |
| 83 | NP_956243.1 | elongation factor 1-beta | eef1b2 | Serine, Threonine, Tyrosine | No | -0.9 | 0.2 | 0.7 | 0.6 | 1.8 | 0.1 | 0.4 | 0.1 | 0.3 | 0.1 |
| 84 | XP_005156056.1 | eukaryotic translation initiation factor 3 subunit C isoform X1 | eif3c | Serine | No | 0.7 | 1.4 | 1.6 | 1.3 | -2.0 | NA | NA | NA | NA | NA |
| 85 | NP_001003763.1 | eukaryotic translation initiation factor 3 subunit H-A | eif3ha | Serine | No | -0.3 | -0.8 | 0.3 | -1.0 | 0.8 | NA | NA | NA | NA | NA |
| 86 | NP_001092707.2 | eukaryotic translation initiation factor 4Ba | eif4ba | Serine, Threonine | YES | 0.9 | 1.0 | 0.9 | 1.2 | 1.2 | 0.0 | 0.1 | 0.1 | 0.2 | 0.1 |
| 87 | XP_005166071.1 | eukaryotic translation initiation factor 4B isoform X1 | eif4bb | Serine, Threonine | YES | 0.7 | 1.2 | 1.1 | 1.1 | 0.2 | 0.0 | 0.1 | 0.1 | 0.1 | 0.1 |
| 88 | XP_005156696.1 | eukaryotic translation initiation factor 4E family member 1c isoform X1 | eif4e1c | Serine, Threonine | No | 0.9 | -0.7 | -0.6 | 0.4 | -1.1 | 0.0 | 0.7 | 1.0 | 0.3 | 0.1 |
| 89 | NP_001007778.1 | eukaryotic translation initiation factor 4eb | eif4eb | Serine | No | 1.2 | 1.0 | 0.8 | 1.2 | 0.8 | NA | NA | NA | NA | NA |
| 90 | XP_021332540.1 | eukaryotic translation initiation factor 4E transporter isoform X1 | eif4enif1 | Serine | YES | 2.0 | 2.2 | 2.0 | 1.9 | 3.4 | NA | NA | NA | NA | NA |
| 91 | XP_021325628.1 | eukaryotic translation initiation factor 4 gamma 3 isoform X1 | EIF4G3 | Serine | YES | 1.0 | 0.7 | 1.2 | 1.3 | 1.8 | NA | NA | NA | NA | NA |
| 92 | NP_001373412.1 | emerin (Emery-Dreifuss muscular dystrophy) isoform 1 | emd | Serine | No | 0.8 | 1.0 | 0.5 | -0.3 | 0.3 | NA | NA | NA | NA | NA |
| 93 | XP_017214412.1 | LOW QUALITY PROTEIN: echinoderm microtubule-associated protein-like 4 | EML4 | Serine, Threonine | No | 0.3 | 0.3 | -0.3 | 0.7 | -1.1 | 0.1 | 0.3 | 0.3 | 0.4 | 0.1 |
| 94 | XP_009292985.1 | protein enabled homolog isoform X1 | enah | Serine | No | 0.6 | 0.2 | -1.4 | -0.1 | -0.2 | NA | NA | NA | NA | NA |
| 95 | XP_686465.3 | epsin-2 isoform X1 | EPN2 | Serine, Threonine, Tyrosine | No | -0.5 | -0.8 | 0.0 | -0.6 | -0.3 | 0.1 | 0.2 | 0.1 | 0.3 | 0.1 |
| 96 | NP_001313413.1 | epiplakin | eppk1 | Serine | No | -0.2 | -0.1 | 0.5 | -0.5 | -0.1 | NA | NA | NA | NA | NA |
| 97 | XP_002663145.3 | epidermal growth factor receptor substrate 15 isoform X1 | EPS15 | Serine | No | -0.3 | -0.7 | 0.1 | -1.0 | 0.4 | NA | NA | NA | NA | NA |
| 98 | XP_005159421.1 | ets2 repressor factor isoform X1 | erf | Serine | YES | 1.1 | 0.8 | 1.5 | 1.2 | 1.1 | NA | NA | NA | NA | NA |
| 99 | XP_005171454.1 | extended synaptotagmin-2-A isoform X1 | ESYT2 | Serine, Threonine | No | 1.3 | 1.8 | 0.9 | 1.3 | 3.0 | 1.1 | 0.8 | 0.8 | 0.7 | 0.9 |
| 100 | XP_005173034.1 | exocyst complex component 7 isoform X10 | exoc7 | Serine, Threonine | YES | 0.4 | 1.4 | 1.1 | 1.4 | 1.0 | 0.0 | 0.1 | 0.1 | 0.2 | 0.1 |
| 101 | NP_957383.2 | exosome component 10 | exosc10 | Serine | No | 0.0 | 0.7 | 0.9 | 0.7 | 0.6 | NA | NA | NA | NA | NA |
| 102 | XP_686077.3 | constitutive coactivator of PPAR-gamma-like protein 2 | FAM120C | Serine, Threonine | No | -0.2 | 0.0 | 0.2 | -0.1 | -0.5 | 0.1 | 0.1 | 0.4 | 0.1 | 0.1 |
| 103 | XP_005170931.1 | protein FAM83G-like isoform X1 | FAM83G | Serine | No | 0.6 | 1.6 | 2.0 | 1.5 | 0.2 | NA | NA | NA | NA | NA |
| 104 | XP_021324662.1 | formin-binding protein 1-like isoform X1 | FNBP1 | Serine, Threonine | No | -0.8 | 0.2 | -0.5 | 0.4 | -1.0 | 0.1 | 0.0 | 0.0 | 0.0 | 0.1 |
| 105 | NP_956196.1 | forkhead box protein K1 | foxk1 | Serine | No | -0.7 | -0.1 | 0.0 | -0.5 | 0.0 | NA | NA | NA | NA | NA |
| 106 | CAP09627.1 | novel protein | fxyd3 | Serine, Threonine | YES | 2.0 | 1.5 | 1.9 | 2.7 | 1.5 | 0.3 | 0.1 | 0.1 | 0.2 | 0.3 |
| 107 | XP_021325450.1 | uncharacterized protein LOC406683 isoform X1 | g3bp2b | Threonine | No | 0.9 | 0.5 | 1.9 | 0.8 | 1.8 | NA | NA | NA | NA | NA |
| 108 | XP_017208591.1 | GRB2-associated-binding protein 1 isoform X1 | gab1 | Serine | No | -0.1 | 0.2 | 0.1 | -0.4 | -0.5 | NA | NA | NA | NA | NA |
| 109 | NP_998259.1 | glyceraldehyde-3-phosphate dehydrogenase 2 | gapdhs | Serine, Threonine | No | -0.6 | -0.2 | 0.6 | -0.6 | 0.5 | 0.0 | 0.1 | 0.1 | 0.2 | 0.1 |
| 110 | XP_009302368.1 | GTPase-activating protein and VPS9 domain-containing protein 1 isoform X1 | gapvd1 | Serine, Threonine | No | -0.2 | -0.3 | -0.4 | 0.3 | -0.4 | 0.0 | 0.1 | 0.1 | 0.2 | 0.1 |
| 111 | XP_009305378.1 | Golgi-specific brefeldin A-resistance guanine nucleotide exchange factor 1 isoform X1 | GBF1 | Serine, Threonine, Tyrosine | No | 0.1 | 0.0 | 0.7 | 0.4 | 0.5 | 0.1 | 0.2 | 0.1 | 0.3 | 0.1 |
| 112 | XP_005155464.1 | G protein-coupled receptor kinase interacting ArfGAP 2 isoform X1 | git2b | Serine, Tyrosine | No | 0.0 | -0.2 | 0.4 | -0.1 | 0.1 | 0.2 | 0.3 | 0.2 | 0.5 | 0.2 |
| 113 | XP_005165358.1 | Golgi phosphoprotein 3 isoform X1 | golph3a | Serine, Tyrosine | YES | 0.5 | 1.2 | 1.4 | 2.2 | 1.4 | 0.2 | 0.3 | 0.2 | 0.5 | 0.2 |
| 114 | NP_957407.1 | homer protein homolog 3b | gpn1 | Threonine | No | -0.6 | 0.4 | 1.6 | 2.1 | 0.1 | NA | NA | NA | NA | NA |
| 115 | NP_571465.1 | glycogen synthase kinase 3 alpha b | gsk3ab | Serine, Threonine, Tyrosine | No | 0.3 | 0.2 | 0.1 | 0.4 | -0.1 | 0.1 | 0.2 | 0.1 | 0.3 | 0.1 |
| 116 | NP_942101.1 | G1 to S phase transition 1, like | gspt1l | Serine | No | 0.1 | 0.3 | 0.2 | 1.1 | 0.4 | NA | NA | NA | NA | NA |
| 117 | NP_001295477.1 | HBS1-like protein | hbs1l | Serine, Threonine | No | 0.2 | 0.3 | 0.3 | 0.1 | 0.6 | 0.0 | 0.1 | 0.1 | 0.2 | 0.1 |
| 118 | XP_005172586.1 | histone deacetylase 4 isoform X1 | hdac4 | Serine, Threonine | No | -0.1 | 0.2 | 0.0 | 0.8 | 0.3 | 0.0 | 0.1 | 0.1 | 0.3 | 0.1 |
| 119 | XP_021335243.1 | heterogeneous nuclear ribonucleoprotein C isoform X1 | hnrnpc | Serine | No | -0.3 | 1.7 | 2.0 | 2.0 | -1.1 | NA | NA | NA | NA | NA |
| 120 | NP_998156.3 | heterogeneous nuclear ribonucleoprotein K, like | hnrpkl | Serine | No | -0.5 | 0.1 | -0.2 | 0.1 | -0.1 | NA | NA | NA | NA | NA |
| 121 | NP_001038538.1 | heat shock protein 90, alpha (cytosolic), class A member 1, tandem duplicate 2 | hsp90aa1.2 | Serine | No | -0.4 | -0.3 | -0.2 | -0.9 | -1.1 | NA | NA | NA | NA | NA |
| 122 | NP_571385.2 | heat shock protein HSP 90-beta | hsp90ab1 | Serine, Threonine | No | 0.2 | 0.8 | 1.0 | 1.3 | 1.1 | 0.1 | 0.2 | 0.2 | 0.2 | 0.3 |
| 123 | XP_009295237.1 | E3 ubiquitin-protein ligase HUWE1 isoform X1 | HUWE1 | Serine, Threonine | No | -0.3 | -0.1 | 1.0 | -0.3 | 0.9 | 0.1 | 0.2 | 0.2 | 0.1 | 0.3 |
| 124 | NP_001034716.1 | cation-independent mannose-6-phosphate receptor precursor | igf2r | Serine | No | 0.0 | -0.2 | 0.0 | 0.4 | 0.3 | NA | NA | NA | NA | NA |
| 125 | NP_956687.1 | immunoglobulin-like domain-containing receptor 1a precursor | ildr1a | Serine, Tyrosine | No | 1.0 | 0.6 | 0.5 | 0.5 | 0.9 | 0.2 | 0.6 | 0.6 | 0.8 | 0.2 |
| 126 | NP_001025236.1 | interleukin enhancer-binding factor 3a | ilf3a | Serine, Threonine | No | -0.4 | -0.5 | -0.1 | 0.0 | -0.1 | 0.0 | 0.1 | 0.0 | 0.2 | 0.1 |
| 127 | NP_997764.1 | interleukin enhancer-binding factor 3 homolog | ilf3b | Threonine | YES | 0.3 | 0.2 | 1.4 | 1.1 | 0.7 | NA | NA | NA | NA | NA |
| 128 | NP_001295895.1 | ras GTPase-activating-like protein IQGAP1 | iqgap1 | Serine, Tyrosine | No | 0.1 | -0.4 | -0.4 | -0.5 | -0.2 | 0.2 | 0.3 | 0.2 | 0.5 | 0.2 |
| 129 | NP_001037819.2 | interferon regulatory factor 2-binding protein 1 | irf2bp1 | Serine | No | 0.3 | -0.1 | 0.2 | -0.3 | 1.3 | NA | NA | NA | NA | NA |
| 130 | NP_999917.1 | interferon regulatory factor 2-binding protein 2-B | irf2bp2b | Serine, Threonine | No | -0.5 | 0.1 | 0.9 | 0.4 | 0.5 | 0.0 | 0.1 | 0.1 | 0.2 | 0.1 |
| 131 | XP_700746.4 | insulin receptor substrate 2 | IRS2 | Threonine | No | 0.4 | -0.4 | 0.6 | 0.5 | 0.7 | NA | NA | NA | NA | NA |
| 132 | XP_005166847.1 | integrin beta-4 isoform X1 | itgb4 | Serine, Threonine, Tyrosine | No | 0.7 | -0.5 | 0.0 | -0.9 | 1.4 | 0.2 | 0.1 | 0.6 | 0.1 | 0.2 |
| 133 | NP_955940.1 | integral membrane protein 2Ba | itm2ba | Threonine | No | -0.3 | -0.2 | 0.7 | 0.3 | 0.8 | NA | NA | NA | NA | NA |
| 134 | XP_005171028.1 | intersectin-1 isoform X1 | ITSN1 | Serine | No | -0.4 | -1.1 | 0.3 | -0.9 | 1.1 | NA | NA | NA | NA | NA |
| 135 | XP_005158835.1 | intersectin-2 isoform X1 | itsn2b | Serine | No | -0.2 | -0.2 | 0.0 | 0.4 | -0.8 | NA | NA | NA | NA | NA |
| 136 | NP_998721.2 | JunB proto-oncogene, AP-1 transcription factor subunit a | junba | Serine | No | 0.6 | 0.7 | 1.3 | -0.4 | 1.3 | NA | NA | NA | NA | NA |
| 137 | NP_001292549.1 | KN motif and ankyrin repeat domain-containing protein 2 | kank2 | Serine | No | 0.2 | -0.3 | -0.7 | -0.5 | -1.1 | NA | NA | NA | NA | NA |
| 138 | XP_009300766.1 | KN motif and ankyrin repeat domain-containing protein 4 isoform X1 | KANK4 | Serine | No | 0.3 | -0.1 | -0.7 | -0.9 | -1.7 | NA | NA | NA | NA | NA |
| 139 | XP_005156690.1 | kinesin light chain 1 isoform X1 | klc1a | Serine | No | -0.5 | 0.3 | 0.5 | 0.6 | -0.3 | NA | NA | NA | NA | NA |
| 140 | XP_005170607.1 | kininogen-1 isoform X1 | kng1 | Serine | YES | 1.1 | 2.6 | 2.6 | 2.1 | 2.5 | NA | NA | NA | NA | NA |
| 141 | NP_998688.2 | keratin, type I cytoskeletal 15 | krt15 | Serine, Threonine, Tyrosine | No | 0.6 | 0.4 | 0.8 | -0.4 | 0.5 | 0.7 | 0.7 | 0.6 | 0.8 | 0.5 |
| 142 | NP_001076574.1 | keratin, type I cytoskeletal 17 | krt17 | Serine, Threonine, Tyrosine | No | -0.5 | 0.4 | 0.4 | 1.6 | 0.0 | 0.5 | 1.4 | 1.1 | 1.0 | 0.3 |
| 143 | NP_848524.1 | keratin, type I cytoskeletal 18 | krt18a.1 | Tyronsine | No | 0.5 | 1.9 | 2.4 | 2.6 | 0.7 | NA | NA | NA | NA | NA |
| 144 | NP_571584.2 | keratin, type II cytoskeletal 4 | krt4 | Serine, Threonine, Tyrosine | No | -0.6 | 0.4 | 1.2 | 0.3 | -0.7 | 0.2 | 0.2 | 0.1 | 0.3 | 0.1 |
| 145 | NP_571231.2 | keratin, type II cytoskeletal 5 | krt5 | Serine, Threonine, Tyrosine | No | 0.1 | -0.3 | -0.1 | -0.3 | -0.1 | 0.4 | 0.3 | 0.3 | 0.3 | 0.1 |
| 146 | NP_001003445.1 | keratin 91 | krt91 | Serine, Threonine, Tyrosine | No | 0.2 | 0.3 | 0.0 | -0.2 | -0.2 | 0.3 | 0.2 | 0.2 | 0.3 | 0.3 |
| 147 | NP_001070922.1 | keratin 94 | krt94 | Serine, Threonine | YES | 0.1 | 1.0 | 3.8 | 3.5 | 3.1 | 0.6 | 0.4 | 0.1 | 0.2 | 0.3 |
| 148 | NP_001002383.1 | keratin 97 | krt97 | Serine, Threonine, Tyrosine | No | -0.2 | -1.1 | -0.7 | -0.7 | 0.0 | 0.1 | 0.3 | 0.2 | 0.1 | 0.1 |
| 149 | XP_021324942.1 | la-related protein 1 isoform X1 | LARP1 | Serine, Threonine | No | 0.6 | 1.0 | 1.5 | 1.6 | 0.3 | 0.1 | 0.1 | 0.1 | 0.0 | 0.0 |
| 150 | NP_571395.2 | plastin-2 | lcp1 | Serine | No | -1.3 | -1.0 | -0.5 | -0.9 | -0.3 | NA | NA | NA | NA | NA |
| 151 | NP_694503.1 | lamin | lmna | Serine, Threonine, Tyrosine | No | -0.3 | -0.5 | -0.4 | -0.1 | 0.1 | 0.1 | 0.3 | 0.3 | 0.2 | 0.3 |
| 152 | NP_571077.2 | lamin-B2 | lmnb2 | Serine | YES | 0.6 | 1.6 | 2.5 | 2.5 | 1.5 | NA | NA | NA | NA | NA |
| 153 | XP_021334325.1 | LIM domain only protein 7 isoform X1 | lmo7a | Serine, Threonine | No | -0.1 | -0.3 | -0.8 | -0.8 | -0.1 | 0.3 | 0.2 | 0.3 | 0.3 | 0.3 |
| 154 | XP_021331684.1 | leucyl-cystinyl aminopeptidase isoform X1 | lnpep | Tyronsine | No | 1.6 | 1.4 | 2.0 | 1.0 | 1.0 | NA | NA | NA | NA | NA |
| 155 | XP_005160562.1 | E3 ubiquitin-protein ligase LNX isoform X1 | lnx1 | Threonine | No | 0.6 | 1.4 | 1.6 | 0.9 | 1.5 | NA | NA | NA | NA | NA |
| 156 | XP_005171950.1 | E3 ubiquitin-protein ligase LRSAM1 isoform X1 | lrsam1 | Serine, Threonine | No | 0.1 | 0.4 | 0.2 | 0.1 | 0.0 | 0.0 | 0.1 | 0.1 | 0.2 | 0.1 |
| 157 | XP_005159030.1 | protein LSM14 homolog A isoform X1 | lsm14ab | Serine | No | -0.8 | -1.2 | 0.1 | -0.7 | 1.0 | NA | NA | NA | NA | NA |
| 158 | NP_001020643.1 | lipolysis-stimulated lipoprotein receptor precursor | lsr | Serine | No | 0.1 | 0.3 | 0.3 | -0.3 | 0.3 | NA | NA | NA | NA | NA |
| 159 | XP_009291414.1 | leucine zipper protein 1 | LUZP1 | Serine | No | 1.3 | 1.3 | 1.5 | 0.2 | 1.9 | NA | NA | NA | NA | NA |
| 160 | NP_001003507.1 | lysM and putative peptidoglycan-binding domain-containing protein 2 | lysmd2 | Serine | No | -0.7 | -1.3 | -0.9 | -0.7 | -0.7 | NA | NA | NA | NA | NA |
| 161 | NP_001032468.2 | dual specificity mitogen-activated protein kinase kinase 2a | map2k2a | Serine, Threonine | No | -0.4 | 0.1 | -0.4 | -0.2 | -0.5 | 0.0 | 0.1 | 0.1 | 0.2 | 0.1 |
| 162 | NP_001121753.1 | dual specificity mitogen-activated protein kinase kinase 2b | map2k2b | Serine, Threonine | YES | -0.9 | -1.0 | -1.0 | -1.1 | -1.5 | 0.0 | 0.1 | 0.1 | 0.2 | 0.1 |
| 163 | XP_021326014.1 | mitogen-activated protein kinase kinase kinase 15 isoform X1 | MAP3K15 | Serine, Threonine | No | -1.3 | -1.1 | -2.1 | -1.7 | -1.3 | 0.0 | 0.1 | 0.1 | 0.2 | 0.1 |
| 164 | NP_001035448.1 | mitogen-activated protein kinase kinase kinase kinase 2 | map4k2 | Serine | No | -0.1 | -0.7 | 0.1 | -1.1 | 0.4 | NA | NA | NA | NA | NA |
| 165 | NP_878308.2 | mitogen-activated protein kinase 1 | mapk1 | Threonine, Tyrosine | YES | 0.1 | 1.1 | 0.9 | 0.5 | 2.0 | 0.6 | 1.2 | 1.1 | 1.1 | 1.6 |
| 166 | CAA75355.1 | stress-activated protein kinase-3 | mapk12a | Threonine, Tyrosine | No | 1.2 | 0.4 | 1.0 | 0.6 | 0.6 | 0.2 | 0.2 | 0.1 | 0.3 | 0.1 |
| 167 | NP_571797.1 | mitogen-activated protein kinase 14A | mapk14a | Threonine, Tyrosine | No | -0.1 | 0.7 | 1.2 | 1.1 | 1.0 | 0.2 | 0.2 | 0.1 | 0.3 | 0.1 |
| 168 | NP_001300688.1 | mitogen-activated protein kinase 14B | mapk14b | Threonine, Tyrosine | No | 0.1 | 0.3 | 0.4 | 0.3 | 0.3 | 0.0 | 0.2 | 0.0 | 0.3 | 0.1 |
| 169 | NP_958915.1 | mitogen-activated protein kinase 3 | mapk3 | Threonine | No | -0.6 | -0.2 | 0.0 | -0.6 | 0.8 | NA | NA | NA | NA | NA |
| 170 | NP_997732.2 | DNA replication licensing factor MCM3 | mcm3 | Serine | YES | 1.3 | 0.8 | 2.1 | 2.0 | 1.2 | NA | NA | NA | NA | NA |
| 171 | NP_001303854.1 | malate dehydrogenase 1Aa, NAD (soluble) isoform mdh1x | mdh1aa | Serine | No | 0.4 | 0.6 | -0.2 | 0.2 | -0.1 | NA | NA | NA | NA | NA |
| 172 | XP_021336132.1 | mediator of RNA polymerase II transcription subunit 1 isoform X1 | MED1 | Serine | No | 0.3 | 0.0 | 0.3 | 0.0 | 0.1 | NA | NA | NA | NA | NA |
| 173 | CAK04954.1 | major histocompatibility complex class I UBA gene | mhc1uba | Serine, Threonine | No | -0.3 | -0.5 | 0.8 | -1.0 | 0.5 | 0.0 | 0.1 | 0.1 | 0.2 | 0.1 |
| 174 | NP_991162.1 | ER membrane protein complex subunit 5 precursor | mmgt1 | Threonine | No | 0.0 | 0.0 | 0.6 | -0.3 | 0.1 | NA | NA | NA | NA | NA |
| 175 | NP_001002518.1 | 39S ribosomal protein L46, mitochondrial | mrpl46 | Tyronsine | No | 1.0 | 1.9 | 1.6 | 2.1 | 1.2 | NA | NA | NA | NA | NA |
| 176 | XP_009302823.1 | E3 ubiquitin-protein ligase MYCBP2 isoform X1 | mycbp2 | Serine | No | 0.0 | -2.1 | -0.5 | -1.6 | 0.5 | NA | NA | NA | NA | NA |
| 177 | XP_009300420.1 | myosin-10 isoform X1 | MYH10 | Serine | No | 0.0 | -0.3 | 0.5 | -0.5 | 0.5 | NA | NA | NA | NA | NA |
| 178 | XP_005165751.1 | myosin-9 isoform X1 | myh9a | Serine, Threonine | No | 0.4 | 0.1 | -0.1 | 0.0 | -0.3 | 0.1 | 0.2 | 0.1 | 0.2 | 0.1 |
| 179 | XP_021322097.1 | unconventional myosin-XVIIIa isoform X1 | MYO18A | Serine, Threonine | No | 0.2 | 1.1 | 1.0 | 1.3 | -0.3 | 0.0 | 0.2 | 0.2 | 0.1 | 0.1 |
| 180 | XP_687465.5 | myoferlin | MYOF | Serine, Threonine | No | 0.0 | 0.3 | 0.5 | 0.2 | 1.1 | 0.1 | 0.3 | 0.2 | 0.3 | 0.3 |
| 181 | NP_997833.2 | nicotinamide phosphoribosyltransferase 2 | nampt2 | Serine | No | 0.7 | -0.2 | 0.4 | -0.2 | 0.6 | NA | NA | NA | NA | NA |
| 182 | XP_009290461.1 | neurobeachin-like protein 2 isoform X1 | NBEAL2 | Serine, Threonine, Tyrosine | No | 0.2 | -0.2 | 0.4 | 0.1 | 0.5 | 0.1 | 0.5 | 0.1 | 0.3 | 0.1 |
| 183 | BAU80756.1 | n-myc downstream-regulated gene 1a-2 | ndrg1a | Serine, Threonine | No | 0.8 | 1.3 | 1.6 | 0.8 | 0.9 | 0.0 | 0.1 | 0.1 | 0.2 | 0.1 |
| 184 | NP_001008593.1 | protein NDRG2 | ndrg2 | Serine, Threonine, Tyrosine | No | -0.3 | -0.4 | -0.4 | -1.2 | -0.2 | 0.1 | 0.4 | 0.3 | 0.4 | 0.1 |
| 185 | XP_005163134.1 | protein NDRG4 isoform X1 | ndrg4 | Serine, Threonine | No | -0.4 | -0.4 | -0.5 | -1.1 | 0.0 | 0.0 | 0.0 | 0.1 | 0.1 | 0.1 |
| 186 | XP_021323721.1 | negative elongation factor E isoform X1 | nelfe | Serine, Threonine | No | -0.4 | -0.2 | -0.2 | 0.1 | -0.7 | 0.0 | 0.1 | 0.1 | 0.2 | 0.1 |
| 187 | NP_001074055.1 | protein Niban 2a | niban2a | Serine, Threonine | YES | 0.2 | 1.0 | 0.8 | 1.0 | 1.2 | 0.0 | 0.1 | 0.2 | 0.2 | 0.1 |
| 188 | NP_001003633.1 | GPN-loop GTPase 1 | nrbp1 | Threonine | No | -0.5 | -0.9 | 0.3 | -0.8 | 0.7 | NA | NA | NA | NA | NA |
| 189 | NP_001038654.2 | nuclear receptor-binding protein | nrbp1 | Threonine | No | -0.4 | -0.5 | -0.1 | -0.3 | -0.2 | NA | NA | NA | NA | NA |
| 190 | NP_001007447.1 | NSFL1 cofactor p47 | nsfl1c | Serine | No | -0.5 | -0.5 | 2.2 | -0.3 | 2.9 | NA | NA | NA | NA | NA |
| 191 | NP_001093536.2 | optineurin | optn | Serine | No | -0.2 | -0.8 | -0.1 | -1.0 | -0.1 | NA | NA | NA | NA | NA |
| 192 | XP_021330869.1 | oxysterol-binding protein-related protein 8 | OSBPL8 | Serine | No | 0.3 | 0.6 | 1.1 | 0.9 | -0.4 | NA | NA | NA | NA | NA |
| 193 | NP_998022.1 | osteoclast-stimulating factor 1 | ostf1 | Serine, Tyrosine | No | 1.0 | 0.5 | 0.0 | 0.9 | 0.2 | 0.2 | 0.1 | 0.0 | 0.3 | 0.2 |
| 194 | NP_957469.2 | serine/threonine-protein kinase OSR1b | oxsr1b | Serine | No | -0.4 | 0.1 | 0.7 | -0.3 | 0.5 | NA | NA | NA | NA | NA |
| 195 | NP_998529.3 | protein disulfide-isomerase precursor | p4hb | Serine | No | -0.1 | -0.4 | -0.2 | -0.9 | 0.0 | NA | NA | NA | NA | NA |
| 196 | NP_956845.1 | phenylalanine-4-hydroxylase | pah | Serine, Threonine, Tyrosine | YES | -0.6 | -0.1 | -1.6 | -1.5 | -1.4 | 0.1 | 0.2 | 0.1 | 0.3 | 0.1 |
| 197 | XP_005171320.1 | serine/threonine-protein kinase PAK 2 isoform X1 | pak2a | Serine, Threonine, Tyrosine | No | -0.3 | 0.1 | 0.1 | -0.1 | 0.6 | 0.1 | 0.2 | 0.1 | 0.3 | 0.1 |
| 198 | XP_021324295.1 | palladin isoform X1 | PALLD | Serine | No | 1.4 | 1.6 | 2.1 | 1.2 | 0.9 | NA | NA | NA | NA | NA |
| 199 | NP_001006015.1 | PRKC apoptosis WT1 regulator protein | pawr | Serine, Threonine | No | 0.7 | 0.5 | 0.3 | -0.6 | 0.1 | 0.0 | 0.1 | 0.1 | 0.2 | 0.1 |
| 200 | NP_998153.1 | programmed cell death protein 4a | pdcd4a | Serine | No | -0.6 | -0.2 | -0.6 | -0.6 | -0.5 | NA | NA | NA | NA | NA |
| 201 | XP_005162743.1 | pyruvate dehydrogenase E1 component subunit alpha, somatic form, mitochondrial isoform X1 | pdha1b | Serine, Tyrosine | No | 0.1 | 1.0 | 1.3 | 0.5 | 1.7 | 0.1 | 0.4 | 0.3 | 0.6 | 0.1 |
| 202 | NP_001313331.1 | PDZ and LIM domain protein 2 | pdlim2 | Serine, Tyrosine | No | 1.8 | 1.9 | 0.5 | 1.1 | 3.5 | 0.1 | 1.0 | 0.8 | 1.1 | 0.1 |
| 203 | NP_956490.1 | PDZ and LIM domain protein 5b isoform 2 | pdlim5b | Serine | No | -0.3 | -3.6 | -3.2 | -1.3 | 0.1 | NA | NA | NA | NA | NA |
| 204 | NP_957302.1 | 6-phosphofructo-2-kinase/fructose-2,6-bisphosphatase 2a | pfkfb2a | Threonine | No | -0.3 | 0.1 | 0.2 | 0.3 | 0.1 | NA | NA | NA | NA | NA |
| 205 | XP_698635.3 | ATP-dependent 6-phosphofructokinase, liver type | PFKL | Serine | No | 0.3 | 0.2 | -0.2 | -1.2 | -1.1 | NA | NA | NA | NA | NA |
| 206 | NP_001315318.1 | phosphofructokinase, liver b | pfklb | Serine, Threonine | No | 0.5 | 0.3 | -0.2 | -1.5 | -0.9 | 0.0 | 0.1 | 0.1 | 0.2 | 0.1 |
| 207 | NP_001007393.1 | membrane-associated progesterone receptor component 1 | pgrmc1 | Threonine, Tyrosine | No | 1.4 | 0.8 | 0.6 | 0.5 | 0.6 | 0.2 | 0.2 | 0.1 | 0.3 | 0.1 |
| 208 | NP_001030149.2 | phosphatidylinositol 4-kinase beta | pi4kb | Serine, Threonine | No | -0.5 | 0.9 | 1.7 | 1.9 | 1.1 | 0.0 | 0.5 | 0.5 | 0.4 | 0.1 |
| 209 | XP_009302996.1 | 1-phosphatidylinositol 3-phosphate 5-kinase isoform X1 | pikfyve | Serine, Threonine | No | 1.0 | 0.7 | 1.0 | 0.4 | 0.4 | 0.0 | 0.1 | 0.1 | 0.2 | 0.1 |
| 210 | XP_005157947.1 | cytoplasmic phosphatidylinositol transfer protein 1 isoform X1 | pitpnc1b | Serine, Threonine | No | 0.5 | 0.6 | 0.2 | -0.3 | -0.1 | 0.1 | 0.2 | 0.1 | 0.2 | 0.1 |
| 211 | NP_001038745.1 | plakophilin-3a | pkp3a | Serine, Threonine | YES | 1.9 | 2.1 | 1.9 | 1.7 | 1.8 | 0.1 | 0.1 | 0.2 | 0.4 | 0.1 |
| 212 | XP_005166590.1 | 1-phosphatidylinositol 4,5-bisphosphate phosphodiesterase beta-3 isoform X1 | plcb3 | Serine | No | 0.0 | 0.0 | -0.2 | -0.2 | 0.2 | NA | NA | NA | NA | NA |
| 213 | NP_001103170.1 | 1-phosphatidylinositol 4,5-bisphosphate phosphodiesterase delta-1a | plcd1a | Serine, Threonine | No | -0.1 | 0.0 | -0.3 | -0.6 | -0.5 | 0.0 | 0.1 | 0.1 | 0.2 | 0.1 |
| 214 | XP_005163413.1 | phospholipase C, delta 1b isoform X1 | plcd1b | Serine | YES | 0.1 | 0.3 | 1.4 | 0.7 | 1.9 | NA | NA | NA | NA | NA |
| 215 | XP_021323775.1 | plectin isoform X1 | pleca | Serine, Threonine, Tyrosine | No | -0.2 | -0.7 | -0.5 | -0.5 | -0.1 | 0.1 | 0.1 | 0.1 | 0.3 | 0.1 |
| 216 | XP_021335504.1 | pleckstrin homology domain-containing family A member 6 isoform X1 | PLEKHA6 | Serine, Tyrosine | No | 1.0 | 3.0 | 1.9 | 1.9 | 1.6 | 0.5 | 0.4 | 0.3 | 0.1 | 0.1 |
| 217 | XP_002662798.2 | pleckstrin homology domain-containing family O member 2 | PLEKHO2 | Serine | No | 0.0 | -0.6 | 0.2 | -0.3 | 1.0 | NA | NA | NA | NA | NA |
| 218 | NP_001074034.1 | uncharacterized protein LOC557029 | plekhs1.3 | Serine | No | -0.5 | -0.4 | -1.0 | -0.7 | -0.3 | NA | NA | NA | NA | NA |
| 219 | XP_009297280.3 | polymerase delta-interacting protein 3 isoform X1 | POLDIP3 | Serine, Threonine | No | 0.5 | 0.6 | 1.0 | 1.0 | 0.4 | 0.0 | 0.1 | 0.1 | 0.2 | 0.1 |
| 220 | XP_005156282.1 | DNA-directed RNA polymerase II subunit RPB1 | POLR2A | Serine, Threonine, Tyrosine | No | -0.1 | 0.1 | 0.3 | 0.9 | -0.1 | 0.1 | 0.2 | 0.2 | 0.2 | 0.1 |
| 221 | XP_005164160.1 | periplakin isoform X1 | ppl | Serine, Threonine | No | -0.5 | -0.1 | 0.4 | -0.6 | 1.9 | 0.1 | 0.3 | 0.3 | 0.3 | 0.4 |
| 222 | XP_021330670.1 | protein phosphatase 1 regulatory subunit 12A isoform X1 | ppp1r12a | Serine, Threonine, Tyrosine | YES | 0.9 | 1.1 | 1.5 | 1.4 | 1.0 | 0.2 | 0.1 | 0.2 | 0.2 | 0.1 |
| 223 | NP_001071047.1 | protein phosphatase 1 regulatory subunit 12C | ppp1r12c | Serine, Threonine | No | -0.9 | 0.8 | 0.8 | 0.7 | -0.1 | 0.5 | 0.4 | 0.5 | 0.4 | 0.2 |
| 224 | NP_001038286.2 | protein phosphatase 2, regulatory subunit B', gamma b isoform 1 | ppp2r5cb | Serine | No | 0.9 | 1.0 | 1.3 | 1.4 | 1.1 | NA | NA | NA | NA | NA |
| 225 | NP_001020680.1 | serine/threonine-protein phosphatase 6 regulatory subunit 2a | ppp6r2a | Serine | No | -0.5 | -0.2 | -0.5 | -0.8 | -0.6 | NA | NA | NA | NA | NA |
| 226 | XP_002663834.2 | PR domain zinc finger protein 2 | PRDM2 | Threonine | No | -2.7 | -2.1 | -3.1 | -3.0 | -3.6 | NA | NA | NA | NA | NA |
| 227 | NP_001009989.1 | protein kinase, cAMP-dependent, regulatory, type I, alpha (tissue specific extinguisher 1) a | prkar1aa | Serine | No | -0.1 | -0.4 | -0.5 | -0.6 | -0.4 | NA | NA | NA | NA | NA |
| 228 | NP_571589.2 | presenilin-2 | psen2 | Threonine | No | -0.2 | 0.1 | 0.4 | -0.2 | 0.7 | NA | NA | NA | NA | NA |
| 229 | NP_001025358.1 | proteasome subunit alpha type-3 | psma3 | Serine | No | 0.2 | 0.0 | 0.0 | 0.3 | -0.1 | NA | NA | NA | NA | NA |
| 230 | NP_001017624.1 | 26S proteasome non-ATPase regulatory subunit 4b | psmd4b | Serine, Threonine | YES | 0.0 | -0.6 | -1.0 | -0.8 | -0.8 | 0.0 | 0.2 | 0.2 | 0.2 | 0.1 |
| 231 | NP_997735.1 | protein tyrosine kinase 2 beta, b | ptk2bb | Serine | No | -0.1 | 0.6 | -0.6 | -0.6 | -1.3 | NA | NA | NA | NA | NA |
| 232 | XP_005161523.1 | tyrosine-protein phosphatase non-receptor type 13 isoform X1 | ptpn13 | Serine, Threonine | No | -0.3 | 0.0 | -0.2 | 0.2 | -0.7 | 0.0 | 0.1 | 0.1 | 0.1 | 0.1 |
| 233 | NP_997819.1 | tyrosine-protein phosphatase non-receptor type 2 | ptpn2b | Serine | YES | 0.4 | 1.5 | 0.9 | 1.1 | 2.3 | NA | NA | NA | NA | NA |
| 234 | XP_687930.5 | tyrosine-protein phosphatase non-receptor type 3 | PTPN3 | Serine | No | -0.7 | -0.2 | -1.0 | -0.6 | -0.8 | NA | NA | NA | NA | NA |
| 235 | XP_021333011.1 | R3H domain-containing protein 2 | R3HDM2 | Serine | No | -0.5 | 1.0 | 1.0 | 0.4 | 0.9 | NA | NA | NA | NA | NA |
| 236 | XP_021326258.1 | guanine nucleotide exchange factor for Rab-3A | RAB3IL1 | Serine | No | -2.2 | -0.5 | -1.3 | -2.4 | -0.1 | NA | NA | NA | NA | NA |
| 237 | NP_571519.1 | guanine nucleotide-binding protein subunit beta-2-like 1 | rack1 | Serine, Threonine | No | -0.5 | -1.1 | -0.6 | -0.8 | -0.9 | 0.0 | 0.1 | 0.1 | 0.2 | 0.1 |
| 238 | XP_017213720.1 | RAF proto-oncogene serine/threonine-protein kinase isoform X1 | raf1b | Serine | No | -0.4 | -0.1 | -1.0 | -0.2 | -1.0 | NA | NA | NA | NA | NA |
| 239 | XP_021322984.1 | ral GTPase-activating protein subunit alpha-1 isoform X1 | RALGAPA1 | Serine | No | -0.3 | 0.0 | -0.4 | -0.1 | -0.1 | NA | NA | NA | NA | NA |
| 240 | XP_021334470.1 | E3 SUMO-protein ligase RanBP2 isoform X1 | RANBP2 | Serine, Threonine | YES | 0.0 | 0.7 | -0.5 | 0.5 | 2.2 | 0.0 | 0.7 | 1.1 | 0.5 | 0.0 |
| 241 | XP_021324455.1 | rap guanine nucleotide exchange factor 1 isoform X1 | rapgef1b | Serine | No | -0.2 | -0.6 | 0.4 | -1.0 | 1.0 | NA | NA | NA | NA | NA |
| 242 | NP_001353656.1 | ras-associated and pleckstrin homology domains-containing protein 1a isoform 1 | RAPH1 | Serine, Threonine | No | 0.3 | 0.7 | 1.0 | 0.8 | -0.6 | 0.0 | 0.1 | 0.1 | 0.2 | 0.1 |
| 243 | XP_005160011.1 | ras-associated and pleckstrin homology domains-containing protein 1 isoform X1 | raph1b | Serine, Threonine | No | -0.1 | 0.8 | 0.7 | 1.3 | 0.1 | 0.0 | 0.2 | 0.2 | 0.1 | 0.1 |
| 244 | XP_009293281.1 | RNA-binding protein 25 isoform X1 | rbm25b | Serine | No | -0.4 | 0.2 | -0.4 | 0.4 | -0.3 | NA | NA | NA | NA | NA |
| 245 | NP_955971.1 | RNA-binding protein 4.2 | rbm4.2 | Serine, Tyrosine | No | 0.6 | 1.3 | 0.6 | 1.7 | 1.3 | 0.3 | 0.4 | 0.6 | 0.1 | 0.2 |
| 246 | XP_005172323.1 | uncharacterized protein LOC406277 isoform X1 | rbm4.3 | Serine | No | 0.2 | 0.3 | 0.1 | 0.7 | 0.3 | NA | NA | NA | NA | NA |
| 247 | NP_001036815.1 | receptor-interacting serine/threonine-protein kinase 1 | ripk1l | Serine | No | 0.1 | 0.3 | 1.1 | 0.4 | 1.2 | NA | NA | NA | NA | NA |
| 248 | NP_001243104.1 | E3 ubiquitin-protein ligase BRE1A | rnf20 | Serine | YES | -1.3 | 0.5 | 0.7 | 0.9 | 0.1 | NA | NA | NA | NA | NA |
| 249 | NP_571655.2 | 60S acidic ribosomal protein P0 | rplp0 | Serine | YES | 1.4 | 2.7 | 2.3 | 3.6 | 2.7 | NA | NA | NA | NA | NA |
| 250 | XP_005162475.1 | regulation of nuclear pre-mRNA domain-containing protein 1B isoform X1 | rprd1b | Serine | No | 0.0 | 0.3 | 0.1 | -0.4 | 0.9 | NA | NA | NA | NA | NA |
| 251 | NP_957447.1 | 40S ribosomal protein S3 | rps3 | Threonine | No | 1.7 | 1.8 | 1.2 | 2.1 | 1.0 | NA | NA | NA | NA | NA |
| 252 | NP_001003728.1 | 40S ribosomal protein S6 | rps6 | Serine, Threonine | No | -1.7 | -1.0 | -0.3 | 2.1 | 0.3 | 0.3 | 0.2 | 0.7 | 0.3 | 0.3 |
| 253 | NP_001139027.1 | ribosome-binding protein 1a | rrbp1a | Serine | YES | -1.1 | -0.3 | -0.4 | 1.5 | 2.3 | NA | NA | NA | NA | NA |
| 254 | NP_001313639.1 | ribosomal L1 domain-containing protein 1 | rsl1d1 | Serine, Threonine | No | 0.0 | -0.2 | -0.4 | -0.8 | -0.8 | 0.0 | 0.1 | 0.1 | 0.2 | 0.1 |
| 255 | XP_005168027.1 | sciellin isoform X1 | scel | Serine, Threonine, Tyrosine | No | 0.0 | 0.1 | -0.2 | 0.4 | 1.3 | 0.3 | 0.5 | 0.5 | 0.4 | 0.3 |
| 256 | XP_021334300.1 | sciellin isoform X2 | scel | Threonine | No | 0.5 | -0.8 | 0.2 | -1.2 | 2.1 | NA | NA | NA | NA | NA |
| 257 | NP_835232.2 | scinderin like a | scinla | Threonine, Tyrosine | YES | -0.9 | -1.7 | -1.1 | -2.5 | -0.6 | 0.2 | 0.2 | 0.1 | 0.3 | 0.1 |
| 258 | NP_998255.2 | scinderin like b | scinlb | Serine, Threonine, Tyrosine | No | -0.1 | 0.5 | -2.0 | -0.7 | -1.0 | 0.8 | 1.4 | 0.5 | 1.4 | 0.7 |
| 259 | XP_009301350.1 | splicing factor 1 isoform X1 | sf1 | Serine | YES | 1.7 | 2.7 | 3.3 | 4.0 | 3.7 | NA | NA | NA | NA | NA |
| 260 | XP_009289295.1 | SH3 domain-containing protein 19 isoform X1 | sh3d19 | Serine | No | -0.1 | 0.2 | 0.7 | 0.4 | 1.2 | NA | NA | NA | NA | NA |
| 261 | XP_005158368.1 | SH3 domain-containing protein 21 isoform X1 | sh3d21 | Threonine | No | 0.6 | 0.8 | 0.3 | -0.1 | -0.4 | NA | NA | NA | NA | NA |
| 262 | AAI71557.1 | Si:xx-by187g17.1 | si:ch211-5k11.8 | Serine, Threonine, Tyrosine | No | -0.4 | -0.9 | 0.5 | -0.5 | -1.0 | 0.1 | 0.2 | 0.1 | 0.3 | 0.1 |
| 263 | XP_009304425.1 | signal-induced proliferation-associated 1-like protein 2 isoform X1 | sipa1l2 | Serine, Threonine | No | -0.1 | 0.5 | 0.3 | 0.6 | -0.1 | 0.0 | 0.1 | 0.1 | 0.2 | 0.1 |
| 264 | NP_998498.2 | SAFB-like transcription modulator | sltm | Serine, Threonine | No | -0.4 | -0.5 | -0.5 | -0.2 | -0.4 | 0.1 | 0.1 | 0.0 | 0.1 | 0.1 |
| 265 | XP_003200294.1 | SWI/SNF complex subunit SMARCC1 isoform X1 | SMARCC1 | Serine | YES | 0.5 | 2.3 | 1.1 | 1.9 | 1.1 | NA | NA | NA | NA | NA |
| 266 | NP_775360.2 | structural maintenance of chromosomes protein 4 | smc4 | Serine | No | 0.8 | 1.7 | -0.5 | 3.1 | 1.8 | NA | NA | NA | NA | NA |
| 267 | NP_001038274.1 | serine/threonine-protein phosphatase 4 regulatory subunit 3 | smek1 | Serine | No | -0.5 | 0.0 | 0.8 | 0.1 | 1.1 | NA | NA | NA | NA | NA |
| 268 | NP_997970.1 | smoothelin, like | smtnl | Serine | YES | 1.7 | 1.1 | 2.4 | 0.8 | 1.4 | NA | NA | NA | NA | NA |
| 269 | NP_001002864.1 | SNW domain-containing protein 1 | snw1 | Serine | YES | 0.4 | 0.9 | 0.8 | 1.1 | 1.1 | NA | NA | NA | NA | NA |
| 270 | NP_991228.1 | sorting nexin-16 | snx16 | Serine | No | 0.1 | -0.2 | 0.7 | -0.2 | 0.4 | NA | NA | NA | NA | NA |
| 271 | XP_009304586.2 | spectrin beta chain, non-erythrocytic 1, partial | SPTBN1 | Serine | No | 0.4 | 0.8 | 0.6 | 0.2 | 1.2 | NA | NA | NA | NA | NA |
| 272 | NP_001093459.1 | spectrin family protein | SPTBN2 | Serine | No | 0.0 | 0.7 | 0.5 | 0.5 | -0.9 | NA | NA | NA | NA | NA |
| 273 | NP_001025246.2 | serine/arginine repetitive matrix protein 2 | srrm2 | Serine | No | 3.5 | 5.4 | 3.2 | 5.5 | 1.4 | NA | NA | NA | NA | NA |
| 274 | AAM34668.1 | arsenite resistance protein 2 | srrt | Serine, Threonine | YES | 0.4 | 1.3 | 1.3 | 1.3 | 0.7 | 0.0 | 0.1 | 0.1 | 0.2 | 0.1 |
| 275 | NP_955870.1 | serine/arginine-rich splicing factor 11 | srsf11 | Serine | No | -0.6 | 1.0 | -0.2 | 1.8 | -0.7 | NA | NA | NA | NA | NA |
| 276 | XP_005157472.1 | serine/arginine-rich splicing factor 1A isoform X1 | srsf1a | Serine, Tyrosine | YES | 1.0 | 2.5 | 1.7 | 2.6 | 0.0 | 0.3 | 0.3 | 0.1 | 0.2 | 0.5 |
| 277 | XP_005172130.1 | lupus La protein isoform X1 | ssb | Serine, Threonine | No | -0.3 | 0.3 | 0.3 | 0.2 | -0.4 | 0.1 | 0.0 | 0.1 | 0.0 | 0.0 |
| 278 | NP_997967.2 | FACT complex subunit SSRP1a | ssrp1a | Tyronsine | No | 0.2 | 1.0 | 0.1 | 1.3 | 0.1 | NA | NA | NA | NA | NA |
| 279 | AAI71583.1 | START domain containing 10 | stard14 | Serine | No | 0.1 | -0.5 | -0.1 | -1.3 | -0.3 | NA | NA | NA | NA | NA |
| 280 | NP_001003984.1 | signal transducer and activator of transcription 5B | stat5b | Serine, Threonine, Tyrosine | No | 0.3 | -0.1 | 0.1 | -0.6 | 0.4 | 0.2 | 0.4 | 0.2 | 0.2 | 0.4 |
| 281 | XP_009294388.1 | striatin-interacting protein 1 homolog isoform X1 | strip1 | Serine | No | -0.1 | -0.7 | -0.1 | -0.4 | 0.0 | NA | NA | NA | NA | NA |
| 282 | NP_001074111.1 | striatin | strn | Serine | No | -0.1 | -0.4 | -0.3 | -0.6 | -0.2 | NA | NA | NA | NA | NA |
| 283 | NP_001003493.1 | sin3 histone deacetylase corepressor complex component SDS3 | suds3 | Threonine | No | 0.1 | 0.8 | 0.1 | 1.0 | -2.2 | NA | NA | NA | NA | NA |
| 284 | XP_005167832.1 | TBC1 domain family member 4 isoform X1 | TBC1D4 | Serine, Threonine | No | -0.4 | 0.0 | 0.1 | -0.3 | 0.1 | 0.3 | 0.1 | 0.0 | 0.2 | 0.0 |
| 285 | NP_956288.1 | transcription elongation factor A protein 1 | tcea1 | Serine, Threonine | No | -0.4 | -0.5 | 0.3 | 0.2 | 0.5 | 0.0 | 0.1 | 0.1 | 0.2 | 0.1 |
| 286 | XP_009301712.1 | transcription factor 12 isoform X4 | tcf12 | Serine | YES | 0.7 | 0.1 | 1.2 | 2.0 | 1.4 | NA | NA | NA | NA | NA |
| 287 | NP_001002721.1 | tuftelin-interacting protein 11 | tfip11 | Serine | YES | 0.7 | 2.0 | 2.5 | 3.1 | 0.9 | NA | NA | NA | NA | NA |
| 288 | NP_999980.1 | thyroid hormone receptor-associated protein 3b | thrap3b | Tyronsine | No | 0.7 | 2.6 | 0.9 | 2.8 | -1.2 | NA | NA | NA | NA | NA |
| 289 | XP_009303467.1 | tight junction protein ZO-2 isoform X1 | tjp2a | Serine, Tyrosine | No | 0.0 | 0.3 | 0.9 | 0.4 | 0.7 | 0.0 | 0.2 | 0.1 | 0.1 | 0.0 |
| 290 | NP_001159390.1 | thymosin beta 1 | tmsb1 | Threonine | No | 0.8 | 1.1 | 0.0 | 0.2 | -0.9 | NA | NA | NA | NA | NA |
| 291 | NP_001254518.1 | 182 kDa tankyrase-1-binding protein | tnks1bp1 | Serine, Threonine | No | 0.4 | 0.1 | 0.4 | -0.3 | 0.6 | 0.0 | 0.1 | 0.1 | 0.2 | 0.1 |
| 292 | XP_021334702.1 | tensin-1 isoform X1 | TNS1 | Serine, Threonine | No | 0.1 | -0.2 | 0.2 | -0.1 | 0.4 | 0.0 | 0.0 | 0.0 | 0.1 | 0.1 |
| 293 | XP_009304960.1 | TOM1-like protein 2 isoform X1 | tom1l2 | Serine | No | -0.5 | -0.2 | -0.9 | -0.5 | -1.4 | NA | NA | NA | NA | NA |
| 294 | XP_021325415.1 | tumor protein D54 isoform X1 | tpd52l2b | Serine | No | 0.1 | -1.1 | -0.2 | -2.0 | -0.7 | NA | NA | NA | NA | NA |
| 295 | NP_956710.1 | transformer-2 protein homolog alpha | tra2a | Serine | No | -0.9 | 1.5 | 1.6 | 2.5 | -1.1 | NA | NA | NA | NA | NA |
| 296 | XP_002663888.2 | LOW QUALITY PROTEIN: trafficking protein particle complex subunit 10 | TRAPPC10 | Serine, Threonine, Tyrosine | No | -0.4 | -0.9 | -0.2 | -0.8 | 0.0 | 0.1 | 0.2 | 0.3 | 0.4 | 0.1 |
| 297 | NP_001002749.1 | tyrosinase-related protein 1b precursor | tyrp1b | Serine, Tyrosine | YES | 0.9 | 0.6 | 2.5 | 2.0 | 3.8 | 0.2 | 0.3 | 0.2 | 0.5 | 0.2 |
| 298 | NP_001186800.1 | ubiquitin carboxyl-terminal hydrolase 3 | usp3 | Serine | No | -0.6 | -0.1 | 0.1 | 0.3 | -0.1 | NA | NA | NA | NA | NA |
| 299 | XP_021323392.1 | ubiquitin carboxyl-terminal hydrolase 8 isoform X1 | USP8 | Serine, Tyrosine | No | 0.8 | 0.6 | 0.5 | 1.0 | 0.2 | 0.2 | 0.3 | 0.2 | 0.5 | 0.2 |
| 300 | XP_021324724.1 | guanine nucleotide exchange factor VAV2 isoform X7 | VAV2 | Threonine | No | 0.2 | 0.0 | 0.0 | 0.2 | 0.3 | NA | NA | NA | NA | NA |
| 301 | NP_001304681.1 | vinculin b | vclb | Threonine | No | -0.4 | -0.1 | -0.2 | -0.4 | 0.0 | NA | NA | NA | NA | NA |
| 302 | XP_005158091.1 | protein virilizer homolog isoform X1 | VIRMA | Serine | No | 0.1 | -0.7 | 0.5 | -1.0 | 1.8 | NA | NA | NA | NA | NA |
| 303 | XP_005157568.1 | vacuolar protein sorting-associated protein 26B isoform X1 | vps26b | Serine, Threonine | YES | -0.5 | -1.9 | -1.1 | -2.2 | -1.2 | 0.0 | 1.9 | 1.8 | 2.0 | 1.6 |
| 304 | NP_001038362.3 | vitellogenin 1 precursor | vtg1 | Serine, Threonine | YES | 2.0 | 3.4 | 1.9 | 5.0 | 0.2 | 0.1 | 0.5 | 0.4 | 0.5 | 0.2 |
| 305 | NP_001038759.2 | vitellogenin 4 precursor | vtg4 | Serine, Threonine | No | 2.5 | 4.1 | 0.0 | 3.9 | 0.8 | 0.2 | 0.4 | 0.4 | 0.2 | 0.5 |
| 306 | XP_001921656.6 | WD repeat-containing protein 26-like | WDR26 | Serine | No | 0.0 | -0.3 | 0.8 | 0.0 | 1.1 | NA | NA | NA | NA | NA |
| 307 | XP_017208486.1 | WD repeat-containing protein 7 | WDR7 | Serine | No | -0.8 | -0.2 | 0.3 | -1.2 | 0.2 | NA | NA | NA | NA | NA |
| 308 | NP_001070115.2 | XIAP-associated factor 1 | xaf1 | Serine | YES | -0.9 | -1.5 | -1.6 | -2.4 | -1.9 | NA | NA | NA | NA | NA |
| 309 | NP_001001944.2 | 5'-3' exoribonuclease 2 | xrn2 | Serine | YES | 0.4 | 1.2 | 1.7 | 1.7 | 2.5 | NA | NA | NA | NA | NA |
| 310 | CAK04259.2 | novel protein similar to vertebrate Yes-associated protein 1 | yap1 | Serine, Threonine | No | 1.2 | 2.5 | 1.1 | 1.1 | 2.7 | 0.0 | 0.1 | 0.1 | 0.2 | 0.1 |
| 311 | AAH86710.1 | Ywhab1 protein, partial | ywhaba | Serine, Threonine, Tyrosine | No | 0.4 | -0.4 | -0.6 | -1.7 | 0.2 | 0.1 | 0.3 | 0.2 | 0.4 | 0.1 |
| 312 | NP_998329.1 | 14-3-3 protein eta | ywhah | Tyronsine | No | -0.3 | -0.8 | -0.5 | -0.8 | 0.2 | NA | NA | NA | NA | NA |
| 313 | NP_958892.1 | tyrosine 3-monooxygenase/tryptophan 5-monooxygenase activation protein, theta polypeptide b | ywhaqb | Serine, Threonine, Tyrosine | No | 0.3 | -0.3 | 0.1 | -1.0 | 0.0 | 0.1 | 0.2 | 0.1 | 0.3 | 0.1 |
| 314 | NP_997922.2 | 14-3-3 protein zeta/delta | ywhaz | Serine, Threonine, Tyrosine | No | 0.1 | -1.3 | -0.9 | -3.2 | -1.2 | 0.1 | 0.4 | 0.3 | 0.3 | 0.1 |
| 315 | NP_956923.1 | zinc finger and BTB domain-containing protein 7B | zbtb7b | Serine, Threonine | No | 0.7 | 0.2 | 0.3 | 0.4 | -0.1 | 0.0 | 0.1 | 0.1 | 0.2 | 0.1 |
| 316 | NP_001020695.2 | zinc finger CCCH domain-containing protein 14 | zc3h14 | Serine | No | -0.9 | -1.1 | -0.7 | -1.3 | -0.6 | NA | NA | NA | NA | NA |
| 317 | NP_001070846.1 | nuclear-interacting partner of ALK | zc3hc1 | Serine | YES | 0.3 | 0.9 | 0.4 | 1.1 | 1.1 | NA | NA | NA | NA | NA |
| 318 | XP_021336562.1 | uncharacterized protein LOC563946 isoform X1 | zgc:136930 | Serine, Threonine, Tyrosine | YES | 0.1 | -0.6 | -1.1 | -2.1 | -1.4 | 0.2 | 0.2 | 0.2 | 0.4 | 0.0 |
| 319 | NP_001083016.1 | LUC7 domain-containing protein | zgc:158803 | Serine, Threonine | No | 0.6 | 1.2 | 1.3 | 1.8 | -1.0 | 0.0 | 0.1 | 0.1 | 0.2 | 0.1 |
| 320 | XP_005166230.1 | uncharacterized protein LOC100136852 isoform X2 | zgc:174863 | Serine | No | 0.7 | -1.0 | -0.6 | -1.0 | 0.0 | NA | NA | NA | NA | NA |
| 321 | XP_017212307.1 | uncharacterized protein LOC327497 isoform X1 | zgc:66473 | Serine | No | -0.1 | 1.1 | -0.2 | 0.3 | 1.1 | NA | NA | NA | NA | NA |
| 322 | XP_009303246.1 | uncharacterized protein LOC445086 isoform X1 | zgc:92380 | Serine, Threonine, Tyrosine | YES | -0.4 | -0.7 | -0.9 | -1.2 | -1.6 | 0.1 | 0.2 | 0.1 | 0.3 | 0.1 |
| 323 | NP_001071004.1 | zinc finger protein 185 | znf185 | Serine, Threonine | No | 0.2 | -0.1 | -0.2 | -0.3 | 0.1 | 0.0 | 0.1 | 0.1 | 0.2 | 0.1 |
| 324 | XP_003201679.5 | coronin-7-like | coro7 | Serine | No | -0.8 | -0.4 | 0.0 | -1.1 | 0.2 | NA | NA | NA | NA | NA |
| 325 | NP_001002649.2 | ATP-citrate synthase | acly | Serine | No | -0.7 | -0.7 | -0.2 | -1.1 | -0.2 | NA | NA | NA | NA | NA |
| 326 | QFX67701.1 | transforming growth factor beta-activated kinase 1b | TAK1 | Serine | YES | -0.7 | -1.0 | -1.2 | -1.0 | -0.8 | NA | NA | NA | NA | NA |
| 327 | XP_009304069.1 | FYN-binding protein isoform X1 | FYB1 | Serine | No | -0.5 | -0.7 | -0.2 | 0.4 | 0.2 | NA | NA | NA | NA | NA |
| 328 | CAQ14849.1 | sortilin 1, like | sort1 | Serine | No | -0.1 | -0.8 | -0.5 | -0.8 | 0.1 | NA | NA | NA | NA | NA |
| 329 | NP_956248.1 | kinesin light chain 1b | klc1 | Serine | No | -0.1 | -0.6 | 1.1 | 0.1 | 0.6 | NA | NA | NA | NA | NA |
| 330 | XP_021330602.1 | non-muscle caldesmon isoform X1 | cald1a | Serine | No | -0.1 | -1.2 | -0.3 | -0.5 | 1.3 | NA | NA | NA | NA | NA |
| 331 | NP_001077019.1 | kinesin light chain 2 | klc2 | Serine | No | 0.2 | -0.4 | 0.8 | -0.2 | 0.7 | NA | NA | NA | NA | NA |
| 332 | Q6PBM7.1 | RecName: Full=Protein FAM76B | FAM76B | Serine | No | 0.2 | -0.1 | 0.5 | -0.1 | 0.2 | NA | NA | NA | NA | NA |
| 333 | XP_003200217.3 | zinc finger CCCH domain-containing protein 13 isoform X1 | zc3h13 | Serine | No | 0.2 | 0.7 | 0.2 | 0.0 | 0.4 | NA | NA | NA | NA | NA |
| 334 | A2RRV3.1 | PAT1-like protein 1 | pat1 | Serine | No | 0.3 | -2.1 | -1.6 | -1.9 | 0.2 | NA | NA | NA | NA | NA |
| 335 | Q502M5.1 | LSM16 protein homolog | LSM16 | Serine | No | 0.3 | 0.2 | 0.9 | 0.7 | 0.3 | NA | NA | NA | NA | NA |
| 336 | XP_021324618.1 | uncharacterized protein LOC110438215 | LOC110438215 | Serine | No | 0.5 | 0.4 | -0.8 | -0.4 | -0.5 | NA | NA | NA | NA | NA |
| 337 | XP_009297834.1 | LIM domain-containing protein 2 isoform X1 | LIMD2 | Serine | No | 1.0 | 1.1 | 0.7 | 0.6 | 0.3 | NA | NA | NA | NA | NA |
| 338 | XP_002663158.1 | uncharacterized protein LOC100330250 | LOC100330250 | Serine | No | 1.2 | -1.2 | -1.4 | -1.2 | 0.4 | NA | NA | NA | NA | NA |
| 339 | XP_003197660.3 | E3 ubiquitin-protein ligase HECW2 | HECW2 | Serine, Threonine | No | -0.9 | -0.6 | -0.3 | -0.3 | -0.1 | 0.0 | 0.1 | 0.1 | 0.2 | 0.1 |
| 340 | XP_017207739.1 | poliovirus receptor-related 2 like isoform X2 | PVRL2 | Serine, Threonine | No | -0.6 | -0.3 | -0.4 | -0.4 | -0.4 | 0.0 | 0.1 | 0.1 | 0.2 | 0.1 |
| 341 | XP_017209059.1 | breast carcinoma-amplified sequence 1 isoform X3 | BCAS1 | Serine, Threonine | No | -0.3 | 0.3 | -0.3 | -1.2 | -0.6 | 0.0 | 0.1 | 0.1 | 0.2 | 0.1 |
| 342 | AAH97213.1 | Krt1-19d protein, partial | KRT1 | Serine, Threonine | No | -0.3 | 0.4 | -0.2 | 0.1 | -0.9 | 0.0 | 0.1 | 0.3 | 0.1 | 0.5 |
| 343 | Q7SXG4.2 | Ubiquitin-like 1-activating enzyme E1B | UBA1 | Serine, Threonine | No | -0.2 | -0.7 | -0.9 | -0.8 | -0.5 | 0.0 | 0.1 | 0.1 | 0.2 | 0.1 |
| 344 | BAN66738.1 | pumilio1 | PUM1 | Serine, Threonine | No | -0.2 | 0.8 | 1.5 | 0.2 | 1.5 | 0.0 | 0.1 | 0.1 | 0.2 | 0.1 |
| 345 | XP_021324257.1 | band 4.1-like protein 2 | Epb41l2 | Serine, Threonine | No | -0.1 | 0.9 | 0.1 | 0.3 | 0.9 | 0.1 | 0.1 | 0.1 | 0.1 | 0.1 |
| 346 | XP_021325750.1 | histone deacetylase 7, partial | HDAC7 | Serine, Threonine | No | -0.1 | -0.8 | 0.0 | -0.5 | 0.5 | 0.0 | 0.4 | 0.3 | 0.3 | 0.1 |
| 347 | Q7ZTS4.2 | Cytokeratin-18 | krt18 | Serine, Threonine | No | 0.0 | 1.0 | 1.7 | 1.6 | 0.5 | 0.3 | 0.3 | 0.3 | 0.2 | 0.0 |
| 348 | NP_001036210.1 | cytoplasmic dynein 1 heavy chain 1 | DYNC1H1 | Serine, Threonine | No | 0.1 | 0.3 | 0.6 | 0.0 | -0.5 | 0.0 | 0.1 | 0.1 | 0.2 | 0.1 |
| 349 | XP_021335365.1 | LIM domain and actin-binding protein 1-like | LIMA1 | Serine, Threonine | YES | 0.4 | 1.6 | 1.5 | 1.8 | 1.5 | 0.5 | 0.7 | 0.4 | 0.9 | 0.3 |
| 350 | A5D8S8.1 | RecName: Full=Protein HEXIM1 | HEXIM1 | Serine, Threonine | No | 0.4 | 1.4 | 1.6 | 1.6 | -0.1 | 0.2 | 0.1 | 0.0 | 0.1 | 0.1 |
| 351 | XP_002661094.3 | myosin-9-like | myh9 | Serine, Threonine | No | 0.5 | 0.2 | 0.7 | 1.0 | 0.5 | 0.1 | 0.1 | 0.1 | 0.2 | 0.1 |
| 352 | NP_001018530.1 | WW domain-binding protein 4 | wbp4 | Serine, Threonine | YES | 0.7 | 1.8 | 1.3 | 1.2 | 1.5 | 0.0 | 0.1 | 0.1 | 0.2 | 0.1 |
| 353 | A0A0R4IKJ1.1 | Phosphorylated CTD-interacting factor 1 | pcif1 | Serine, Threonine | No | 0.7 | 2.6 | 0.4 | 1.3 | 2.6 | 0.0 | 0.1 | 0.1 | 0.2 | 0.1 |
| 354 | NP_001243566.1 | desmoglein-2.1 precursor | dsg2 | Serine, Threonine | No | 0.9 | -0.9 | -0.2 | -0.2 | 0.4 | 0.0 | 0.2 | 0.1 | 0.2 | 0.1 |
| 355 | XP_002661360.3 | proline and serine-rich protein 2 | PROSER2 | Serine, Threonine | No | 0.9 | 1.8 | 0.5 | 0.5 | 2.4 | 0.4 | 1.0 | 0.4 | 0.5 | 0.6 |
| 356 | XP_005173887.2 | desmoplakin-like | dsp | Serine, Threonine | No | 0.9 | 0.6 | 0.4 | -1.0 | 0.3 | 0.0 | 0.1 | 0.1 | 0.2 | 0.1 |
| 357 | AAI54493.1 | Rps6ka1 protein, partial | Rps6ka1 | Serine, Threonine | No | 1.2 | 3.0 | -0.1 | 3.0 | 0.4 | 0.4 | 0.4 | 0.6 | 0.3 | 1.3 |
| 358 | XP_021326591.1 | vitellogenin-like | vtg | Serine, Threonine | YES | 1.8 | 2.9 | 2.4 | 5.1 | 1.5 | 0.3 | 0.7 | 0.7 | 0.6 | 0.1 |
| 359 | AAI16561.1 | Sept2 protein, partial | Sept2 | Serine, Threonine, Tyrosine | No | -1.7 | -0.1 | 0.4 | -0.5 | 2.1 | 0.1 | 0.2 | 0.1 | 0.3 | 0.1 |
| 360 | NP_571106.2 | actin, cytoplasmic 1 | actb1 | Serine, Threonine, Tyrosine | No | -1.4 | -1.5 | -1.6 | -1.6 | -1.2 | 0.1 | 0.2 | 0.1 | 0.3 | 0.1 |
| 361 | NP_001035074.1 | eukaryotic translation elongation factor 1 alpha 1, like 2 | eEF1A1 | Serine, Threonine, Tyrosine | No | -1.3 | -1.9 | -0.3 | -1.7 | 0.1 | 0.1 | 0.2 | 0.1 | 0.3 | 0.1 |
| 362 | AAB03704.1 | heat shock cognate | hsp70 | Serine, Threonine, Tyrosine | No | -0.5 | 0.0 | 0.8 | -0.5 | 1.1 | 0.1 | 0.2 | 0.1 | 0.3 | 0.1 |
| 363 | XP_009301181.2 | uncharacterized protein si:ch73-368j24.3 | si:ch73-368j24.3 | Serine, Threonine, Tyrosine | No | -0.3 | -0.6 | -1.2 | -1.2 | -0.9 | 0.1 | 0.2 | 0.1 | 0.3 | 0.2 |
| 364 | XP_021325562.1 | plakophilin-1 | PKP1 | Serine, Threonine, Tyrosine | No | -0.1 | -0.6 | -0.6 | -0.7 | -0.4 | 0.2 | 0.1 | 0.2 | 0.1 | 0.1 |
| 365 | XP_021333014.1 | sciellin-like | scel | Serine, Threonine, Tyrosine | No | 0.2 | -0.2 | -0.6 | -0.2 | 0.1 | 0.1 | 0.2 | 0.1 | 0.3 | 0.2 |
| 366 | Q6NWF6.1 | Cytokeratin-8 | krt8 | Serine, Threonine, Tyrosine | No | 0.3 | 0.3 | 0.2 | 0.4 | -0.7 | 0.4 | 0.4 | 0.5 | 0.7 | 0.5 |
| 367 | Q6PE18.1 | Phosphatidylinositol 4-kinase type 2-alpha | pi4k2a | Serine, Tyrosine | No | 0.1 | 0.1 | 0.1 | 0.1 | 0.0 | 0.2 | 0.3 | 0.2 | 0.5 | 0.2 |
| 368 | B5DE31.1 | Y-box-binding protein 1 | ybx1 | Serine, Tyrosine | YES | 0.8 | 1.1 | 2.0 | 1.8 | 1.6 | 0.2 | 0.3 | 0.2 | 0.5 | 0.2 |
| 369 | AAI51865.1 | Wu:fb15e04 protein, partial | Wu:fb15e04 | Threonine | No | -0.2 | -0.1 | -1.6 | -1.4 | -2.0 | NA | NA | NA | NA | NA |
| 370 | XP_005164409.1 | uncharacterized protein LOC101884073 | LOC101884073 | Threonine | No | -0.2 | 0.1 | 1.2 | -0.8 | 1.9 | NA | NA | NA | NA | NA |
| 371 | XP_021322409.1 | oxidation resistance protein 1 isoform X1 | OXR1 | Threonine | No | 0.4 | 0.8 | 0.5 | -0.3 | 0.6 | NA | NA | NA | NA | NA |
| 372 | XP_017209912.1 | probable ATP-dependent RNA helicase DDX41 | DDX41 | Threonine | No | 1.4 | 1.8 | 2.7 | 2.9 | 1.7 | NA | NA | NA | NA | NA |
| 373 | XP_021326418.1 | ras and EF-hand domain-containing protein-like | RASEF | Tyronsine | No | -0.2 | 0.0 | -0.3 | 0.1 | 0.8 | NA | NA | NA | NA | NA |
| 374 | AAI35069.1 | Krt1-19d protein, partial | krt1 | Tyronsine | No | 0.2 | 0.9 | -0.8 | 0.5 | -0.9 | NA | NA | NA | NA | NA |
| 375 | XP_021325269.1 | receptor-type tyrosine-protein phosphatase C isoform X1 | ptprc | Serine | No | -1.4 | 0.0 | -0.2 | -0.8 | 0.2 | NA | NA | NA | NA | NA |
| 376 | XP_005170404.1 | msx2-interacting protein isoform X1 | spen | Serine | No | -1.4 | -0.4 | 0.2 | -0.1 | 0.0 | NA | NA | NA | NA | NA |
| 377 | XP_005171185.1 | plasminogen activator inhibitor 1 RNA-binding protein isoform X1 | SERBP1 | Serine | No | -1.3 | -0.5 | -0.3 | 0.0 | -0.5 | NA | NA | NA | NA | NA |
| 378 | XP_009292755.1 | uncharacterized protein prdm1b | prdm1b | Serine | No | -1.2 | -1.9 | -0.8 | -2.3 | -0.5 | NA | NA | NA | NA | NA |
| 379 | XP_005170208.1 | ubiquitin carboxyl-terminal hydrolase 24 isoform X1 | USP24 | Serine | No | -0.6 | -0.2 | -0.5 | -0.1 | -0.5 | NA | NA | NA | NA | NA |
| 380 | XP_009293997.1 | protein NLRC3 | NLRC3 | Serine | YES | -0.6 | -0.9 | -1.1 | -0.5 | -0.8 | NA | NA | NA | NA | NA |
| 381 | XP_021333430.1 | zinc finger protein 638 isoform X2 | Zfp638 | Serine | No | -0.6 | -0.1 | 0.3 | 0.0 | -2.2 | NA | NA | NA | NA | NA |
| 382 | XP_005173279.1 | rho guanine nucleotide exchange factor 6 isoform X1 | ARHGEF6 | Serine | No | -0.5 | -0.6 | 0.1 | -0.6 | 0.2 | NA | NA | NA | NA | NA |
| 383 | XP_009303774.1 | C2 domain-containing protein 2 isoform X1 | C2CD2 | Serine | No | -0.3 | 0.1 | 0.3 | 0.6 | 0.4 | NA | NA | NA | NA | NA |
| 384 | XP_005162978.2 | microtubule-associated protein 4-like isoform X1 | MAP4 | Serine | No | -0.3 | -0.2 | -0.5 | -0.8 | -0.5 | NA | NA | NA | NA | NA |
| 385 | XP_021325325.1 | arf-GAP with coiled-coil, ANK repeat and PH domain-containing protein 2-like | ACAP2 | Serine | No | -0.3 | 0.4 | 0.3 | 0.1 | 0.8 | NA | NA | NA | NA | NA |
| 386 | Q90XF2.2 | Protein kinase C iota type | prkci | Serine | YES | -0.3 | 1.2 | 2.2 | 1.2 | 2.4 | NA | NA | NA | NA | NA |
| 387 | XP_690835.5 | kanadaptin isoform X1 | Slc4a1ap | Serine | No | -0.2 | 0.1 | 0.2 | -0.4 | 0.0 | NA | NA | NA | NA | NA |
| 388 | XP_005174598.3 | caskin-2-like isoform X1 | CASKIN2 | Serine | No | -0.2 | 0.2 | 0.5 | -0.4 | 0.0 | NA | NA | NA | NA | NA |
| 389 | XP_699731.6 | extended synaptotagmin-1 isoform X1 | ESYT1 | Serine | No | -0.2 | -0.9 | -0.6 | -1.2 | -0.7 | NA | NA | NA | NA | NA |
| 390 | XP_005155592.1 | zinc finger FYVE domain-containing protein 16 isoform X1 | ZFYVE16 | Serine | No | -0.2 | -0.4 | 0.0 | -0.5 | 0.9 | NA | NA | NA | NA | NA |
| 391 | XP_005173736.1 | vasodilator-stimulated phosphoprotein isoform X1 | VASP | Serine | No | -0.2 | 0.2 | 0.0 | 0.1 | -0.2 | NA | NA | NA | NA | NA |
| 392 | XP_021330865.1 | mucin-5AC | MUC5AC | Serine | No | -0.1 | 0.2 | -0.1 | -0.1 | 0.1 | NA | NA | NA | NA | NA |
| 393 | XP_009293975.2 | NACHT, LRR and PYD domains-containing protein 3-like | Nlrp3 | Serine | YES | 0.1 | -2.2 | -2.0 | -0.9 | -0.5 | NA | NA | NA | NA | NA |
| 394 | XP_021330036.1 | neuroblast differentiation-associated protein AHNAK isoform X1 | AHNAK | Serine | No | 0.1 | -2.3 | -1.4 | -2.3 | 0.6 | NA | NA | NA | NA | NA |
| 395 | XP_021334691.1 | sperm-specific antigen 2 | ITPRID2 | Serine | No | 0.4 | -1.5 | -1.0 | -0.1 | -0.5 | NA | NA | NA | NA | NA |
| 396 | XP_005173569.1 | protein capicua homolog isoform X1 | CIC | Serine | No | 0.4 | 0.4 | 0.7 | 0.6 | 0.1 | NA | NA | NA | NA | NA |
| 397 | XP_701004.4 | GAS2-like protein 1 | GAS2 | Serine | No | 0.6 | -0.2 | 0.7 | 0.0 | 1.0 | NA | NA | NA | NA | NA |
| 398 | XP_002664782.2 | protein IWS1 homolog isoform X1 | IWS1 | Serine | No | 0.6 | 2.0 | 1.5 | 2.4 | -1.6 | NA | NA | NA | NA | NA |
| 399 | XP_021328549.1 | protein phosphatase Slingshot homolog 3 | SSH3 | Serine | No | 0.6 | 1.0 | 0.7 | 0.1 | -0.2 | NA | NA | NA | NA | NA |
| 400 | XP_005158614.1 | la-related protein 1B isoform X1 | LARP1B | Serine | No | 0.9 | 0.2 | 0.2 | 0.2 | 0.4 | NA | NA | NA | NA | NA |
| 401 | XP_021329227.1 | electromotor neuron-associated protein 1 | SMN1 | Serine | No | 1.0 | 4.1 | -3.4 | 3.3 | 3.1 | NA | NA | NA | NA | NA |
| 402 | XP_017211889.1 | nuclear pore complex protein Nup214 | Nup214 | Serine, Threonine | No | -0.8 | -0.6 | 0.5 | 0.9 | 0.9 | 0.0 | 0.1 | 0.1 | 0.2 | 0.1 |
| 403 | NP_001373713.1 | zinc finger BED domain-containing protein isoform 1 | Zf-BED | Serine, Threonine | No | -0.8 | -0.9 | -0.3 | -0.8 | 0.1 | 0.2 | 0.3 | 0.2 | 0.3 | 0.1 |
| 404 | XP_009302595.1 | FK506-binding protein 15 isoform X1 | FKBP15 | Serine, Threonine | No | -0.3 | 0.0 | 0.4 | 0.2 | 0.1 | 0.1 | 0.2 | 0.0 | 0.2 | 0.2 |
| 405 | XP_684861.3 | absent in melanoma 1 protein isoform X1 | AIM1L | Serine, Threonine | No | -0.1 | 0.6 | 0.8 | 0.0 | 0.6 | 0.2 | 0.2 | 0.2 | 0.2 | 0.0 |
| 406 | XP_021336807.1 | LIM and calponin homology domains-containing protein 1 isoform X1 | LIMCH1 | Serine, Threonine | No | 0.1 | 0.0 | -1.1 | -0.6 | 0.1 | 0.0 | 0.3 | 0.4 | 0.4 | 0.1 |
| 407 | XP_005173195.1 | lysine-specific demethylase 3B isoform X1 | KDM3B | Serine, Threonine | No | 0.1 | 0.4 | 0.7 | 1.2 | 0.6 | 0.0 | 0.2 | 0.1 | 0.2 | 0.1 |
| 408 | XP_001333900.3 | uncharacterized protein si:ch211-288g17.4 | si:ch211-288g17.4 | Serine, Threonine | No | 0.3 | 0.2 | 0.0 | 0.6 | -0.5 | 0.0 | 0.1 | 0.1 | 0.2 | 0.1 |
| 409 | XP_002666105.5 | protein POF1B | POF1B | Serine, Threonine | No | 0.4 | -1.0 | -0.2 | -0.1 | -0.3 | 0.1 | 0.0 | 0.0 | 0.1 | 0.3 |
| 410 | XP_021328593.1 | synergin gamma isoform X1 | SYNRG | Serine, Threonine | No | 0.7 | 0.7 | 1.2 | 0.7 | 0.8 | 0.0 | 0.1 | 0.1 | 0.1 | 0.1 |
| 411 | XP_009304816.1 | supervillin isoform X1 | SVIL | Serine, Threonine | YES | 0.7 | 0.9 | 1.3 | 1.1 | 1.2 | 0.2 | 0.5 | 0.5 | 0.7 | 0.2 |
| 412 | XP_001338671.2 | caldesmon, smooth muscle-like isoform X1 | CALD1 | Serine, Threonine | YES | 0.7 | 1.5 | 2.2 | 2.2 | 0.7 | 0.6 | 0.1 | 0.1 | 0.2 | 0.9 |
| 413 | XP_021324336.1 | uridine-cytidine kinase-like 1 isoform X1 | UCKL1 | Serine, Threonine | YES | 2.5 | 4.1 | 4.4 | 4.4 | 2.7 | 0.7 | 0.1 | 0.1 | 0.2 | 0.8 |
| 414 | XP_009304444.1 | collagen alpha-1(VI) chain | COL6A1 | Serine, Threonine, Tyrosine | No | -1.1 | -2.1 | -1.8 | -2.6 | -1.2 | 0.2 | 0.6 | 0.5 | 0.6 | 0.2 |
| 415 | XP_021334590.1 | uncharacterized protein col6a3 isoform X1 | COL6A3 | Serine, Threonine, Tyrosine | YES | -1.0 | -0.7 | -1.3 | -1.1 | -1.7 | 0.1 | 0.2 | 0.1 | 0.3 | 0.1 |
| 416 | Q6PC29.1 | RecName: Full=14-3-3 protein gamma-1 | YWHAG | Serine, Threonine, Tyrosine | No | -0.5 | -0.5 | -0.3 | -1.7 | -0.2 | 0.1 | 0.2 | 0.1 | 0.3 | 0.1 |
| 417 | XP_691943.2 | histone H2B 1/2-like | H2B | Serine, Threonine, Tyrosine | No | 0.0 | -0.9 | -1.7 | -1.8 | -1.3 | 0.5 | 0.6 | 0.1 | 0.4 | 0.3 |
| 418 | XP_005172514.1 | uncharacterized protein LOC393431 isoform X1 | LOC393431 | Serine, Threonine, Tyrosine | YES | 0.0 | -0.7 | -1.7 | -1.9 | -1.5 | 0.1 | 0.2 | 0.1 | 0.3 | 0.1 |
| 419 | XP_021322489.1 | plectin isoform X1 | PLEC | Serine, Threonine, Tyrosine | No | 0.1 | -0.5 | -0.5 | -0.6 | 0.0 | 0.1 | 0.2 | 0.2 | 0.3 | 0.1 |
| 420 | XP_005173239.1 | neuroblast differentiation-associated protein AHNAK isoform X1 | AHNAK | Serine, Threonine, Tyrosine | No | 0.3 | 0.3 | 0.2 | -0.3 | 0.2 | 0.2 | 0.1 | 0.2 | 0.2 | 0.2 |
| 421 | XP_005165493.1 | drebrin isoform X1 | DBN1 | Serine, Threonine, Tyrosine | No | 0.4 | 0.1 | 0.5 | 0.5 | 0.5 | 0.3 | 0.3 | 0.2 | 0.3 | 0.1 |
| 422 | XP_697241.4 | collagen alpha-1(XVII) chain isoform X1 | COL17A1 | Serine, Threonine, Tyrosine | No | 0.5 | 0.2 | 0.0 | 0.1 | 0.6 | 0.1 | 0.2 | 0.2 | 0.3 | 0.1 |
| 423 | XP_005164248.1 | calponin homology domain-containing protein DDB_G0272472 isoform X1 | LRCH1 | Serine, Threonine, Tyrosine | YES | 1.7 | 2.0 | 2.3 | 2.0 | 1.3 | 0.1 | 0.2 | 0.1 | 0.3 | 0.1 |
| 424 | XP_005165804.1 | disabled homolog 2-interacting protein | DAB2 | Serine, Tyrosine | YES | -1.7 | -1.0 | -0.8 | -0.3 | -1.1 | 0.2 | 0.4 | 0.4 | 0.2 | 0.2 |
| 425 | XP_002666222.2 | uncharacterized protein wu:fi37e09 | wu:fi37e09 | Serine, Tyrosine | No | -1.3 | -1.2 | -2.2 | -2.1 | -1.8 | 0.2 | 0.3 | 0.2 | 0.5 | 0.2 |
| 426 | Q6NYE2.1 | RecName: Full=Protein RCC2 homolog | RCC2 | Serine, Tyrosine | No | -0.7 | -1.0 | 0.2 | -0.5 | 0.1 | 0.2 | 0.3 | 0.2 | 0.5 | 0.2 |
| 427 | XP_005173179.1 | matrin 3-like isoform X1 | MATR3 | Threonine | No | -1.9 | 0.3 | 1.0 | -0.5 | 1.3 | NA | NA | NA | NA | NA |
| 428 | XP_017211485.1 | sorbin and SH3 domain-containing protein 2 isoform X1 | SORBS2 | Threonine | No | -0.1 | 0.1 | -0.4 | -0.3 | 1.1 | NA | NA | NA | NA | NA |
| 429 | XP_002663914.4 | growth arrest-specific protein 7-like | GAS7 | Threonine | No | 0.1 | 0.2 | 0.0 | 0.2 | -0.2 | NA | NA | NA | NA | NA |
| 430 | XP_021325949.1 | sickle tail protein homolog isoform X1 | Skt | Threonine | No | 0.1 | 0.3 | 0.1 | 0.7 | 0.4 | NA | NA | NA | NA | NA |
| 431 | XP_009305121.1 | brain-specific angiogenesis inhibitor 1-associated protein 2 isoform X1 | Baiap2 | Threonine | No | 0.2 | 0.2 | 0.3 | -0.5 | 0.0 | NA | NA | NA | NA | NA |
| 432 | XP_700212.1 | E3 ubiquitin/ISG15 ligase TRIM25 | TRIM25 | Threonine | No | 0.2 | 1.4 | 0.6 | 0.7 | 1.2 | NA | NA | NA | NA | NA |
| 433 | XP_695258.2 | zinc finger protein 839 isoform X1 | ZNF839 | Threonine | No | 0.9 | 1.3 | 0.4 | 0.6 | -0.6 | NA | NA | NA | NA | NA |
| 434 | XP_021335989.1 | tripartite motif-containing protein 14-like | TRIM14 | Threonine | No | 1.1 | 0.6 | 0.5 | 1.3 | 1.9 | NA | NA | NA | NA | NA |
| 435 | XP_687624.2 | rho guanine nucleotide exchange factor 33 isoform X2 | Arhgef33 | Threonine | No | 1.1 | 0.6 | 0.5 | 1.3 | 1.9 | NA | NA | NA | NA | NA |
| 436 | Q1LVV0.1 | Protein FAM83H | FAM83H | Threonine | No | 1.1 | 1.6 | 2.2 | -1.0 | 3.0 | NA | NA | NA | NA | NA |
| 437 | XP_003201508.2 | non-muscle caldesmon-like | CALD1 | Threonine | No | 2.8 | 2.2 | 1.5 | 2.4 | 1.3 | NA | NA | NA | NA | NA |
| 438 | XP_005170784.2 | histone H1-like | H1 | Threonine, Tyrosine | No | -1.2 | -0.3 | -0.7 | -0.4 | -1.5 | 0.2 | 0.2 | 0.1 | 0.3 | 0.1 |
| 439 | AAH92980.1 | Si:dkeyp-113d7.4 protein, partial | Si:dkeyp-113d7.4 | Threonine, Tyrosine | No | 0.0 | -0.2 | -1.1 | -0.8 | -0.7 | 1.0 | 1.5 | 0.6 | 1.7 | 0.0 |
| 440 | AAP37456.1 | signal transducer and activator of transcription 5 | STAT5 | Tyronsine | No | 0.9 | 0.8 | 0.8 | 0.9 | 0.7 | NA | NA | NA | NA | NA |
